# A microproteome screen identifies immunomodulatory bacterial microproteins encoded in expanded gene arrays in *Leptotrichia*

**DOI:** 10.64898/2026.08.11.744246

**Authors:** Mark Ragheb, Yuya Kiguchi, Jordan D. Lin, Florian T. Hoffman, Leighton Daigh, Meenakshi Chakraborty, Boryana Doyle, Matthew P. Grieshop, Ann Lin, Dylan Maghini, Kaitlyn Spees, Lacramioara Bintu, Michael C. Bassik, Ami S. Bhatt

## Abstract

The human microbiome exerts broad influence in health and disease with associative studies implicating the microbiome in influencing immunity, cancer outcomes, and neurodegeneration. However, the molecular mediators of microbe-host communication remain poorly defined. Bacterial microproteins from the microbiome represent a largely uncharacterized class of potential regulators of host immunity. Here, we utilize functional genomics to interrogate 3,552 microproteins in order to identify novel microbial-immune interactions. We constructed a microproteome library from microbial metagenomic datasets, expressed it in macrophages and assayed for immunomodulatory activity. We identify several bacterial microproteins that drive macrophage M1 polarization. Among the strongest hits are a cluster of structurally related microproteins from *Leptotrichia* species, which are oral Gram-negative commensals associated with differential cancer outcomes. Genomic analysis reveals that *Leptotrichia* species encode these putative immunomodulatory microproteins in tandem arrays of up to 44 copies. These genes encode microproteins with varying sequences but conserved predicted structures. In an orthogonal approach, we demonstrate that bacterial expression of *Leptotrichia* microproteins influences macrophage cell state and function. As a whole, our findings identify novel microbial microproteins with immunomodulatory activity and provide a framework for future discovery of host-microbe interactions that influence human health.

## Introduction

Diverse mechanisms of crosstalk have evolved over millennia to facilitate the delicate relationship between the microbiome and the human host. The immune system is often at the front line of this communication, mediating recognition, tolerance, and organismal homeostasis at the microbe-host interface^1–5^. Disruptions of these interactions are implicated in a wide range of disease states, including autoimmunity^6,7^, neurodegeneration^8,9^, and cancer^10^. For example, gut commensal bacteria such as *Candidatus arthromitus* (commonly known as segmented filamentous bacteria) drive intestinal Th17 cell accumulation and IL-17 production, with subsequent implications for inflammatory bowel disease and for systemic autoimmunity^11–13^. Multiple commensal microbes have also been linked to differential cancer outcomes. For example, in addition to driving transformation^14^, *Fusobacterium nucleatum* also facilitates colorectal cancer immune evasion through various proposed mechanisms, including macrophage polarization^15^. Commensal bacteria also shape anti-tumor immune responses, as the microbiome composition predicts anti-PD-1 immunotherapy response in melanoma patients, and fecal microbiota transfer from responders can reprogram the tumor immune microenvironment in refractory disease^16–18^. While the number of microbiome disease associations continues to expand, identifying the underlying molecular interactions that drive these associations remains a significant challenge. An interest in mechanistically cataloging the vast chemical and molecular space of potential microbiome-host interactions has led to systematic screening approaches. For example, unbiased screens have identified commensal bacteria that produce N-acyl amide GPCR ligands that mimic host signalling molecules^19,20^, and proteome-scale interaction mapping has revealed a large number of direct microbiome- exoprotein interactions, including strains of *Fusobacterium* which bind to the inhibitory receptor SIRPɑ and reduce macrophage phagocytosis^21^.

Of the various compounds that mediate microbe-host communication, microbial small molecules and secondary metabolites have been the most extensively studied. For example, butyrate drives regulatory T cell induction via histone modifications^22,23^, and secondary bile acids regulate both innate and adaptive immune cell fate via signaling through cell surface and nuclear receptors^23,24^. Microbial microproteins represent intriguing and largely unexplored candidates for microbe-host communication. Classically defined as proteins of 50 amino acids or fewer^25^, microproteins represent a vast and largely unexplored fraction of the bacterial proteome, due to challenges in prediction and annotation methods. While computational analyses of metagenomic data have recently unearthed thousands of new predicted microproteins^25–28^, functional characterization remains lagging. Nonetheless bacterial microproteins are known to mediate extracellular microbial signalling interactions, such as in quorum sensing^29^ and virulence responses^30^. Interestingly, computationally identified microproteins often contain predicted signal peptides that enable surface expression or secretion, which could facilitate direct contact with host cells^25^. Efforts to de-orphan recently annotated microbial microproteins have led to the discovery of novel antimicrobial peptides^31–33^, yet systematic approaches to identify microproteins that modulate host immune cells remains limited^34^.

In this work, we use a peptide display platform to perform functional genomic screens, assaying a library of over 3000 microproteins for immunomodulatory activity in macrophages. We identify multiple pro-inflammatory microproteins, including a family of structurally related microproteins from *Leptotrichia* species that are uniquely encoded in tandem genomic arrays. Interrogation of these related *Leptotrichia* microproteins reveals that they promote an inflammatory macrophage cell state, and enhance phagocytic activity. These findings identify a previously unrecognized class of putative microbial immunomodulators and establish a scalable framework for systematic discovery of microprotein mediated microbiome-immune interactions.

## Results

### Construction of a bacterial microprotein library to screen for immunomodulatory activity

To identify microproteins that could plausibly signal to host immune cells, we mined multiple metagenomic datasets, searching over 2,000 bacterial metagenomes. We sourced the majority of metagenomes from the NIH Human Microbiome Project (HMP-II)^25,35^ and to enrich for microproteins with putative immune activity, we also included metagenomic samples from patients with inflammatory bowel disease (IBD)^36^ and patients with differential response to immunotherapy^16^. In addition to microbial microproteins identified from metagenomes, we included candidate human G-protein coupled receptor (GPCR) ligands including known formyl peptide receptor ligands^37^. In total, we identified over two million bacterial microproteins to further analyze (Fig. 1a). To prioritize microproteins most likely to have immunomodulatory activity, we filtered for microproteins with the presence of signal peptides using SignalP-6.0^38^, reasoning that secreted or membrane-associated microproteins are best positioned for cell-cell communication. Sequences were dereplicated based on amino acid homology, and ambiguous amino acids (i.e. any codon with an unknown or degenerate base) were converted to glycine, yielding a final list of 3,552 microproteins for downstream screening (Fig. 1a, Supplementary Table 1).

**Figure 1.**
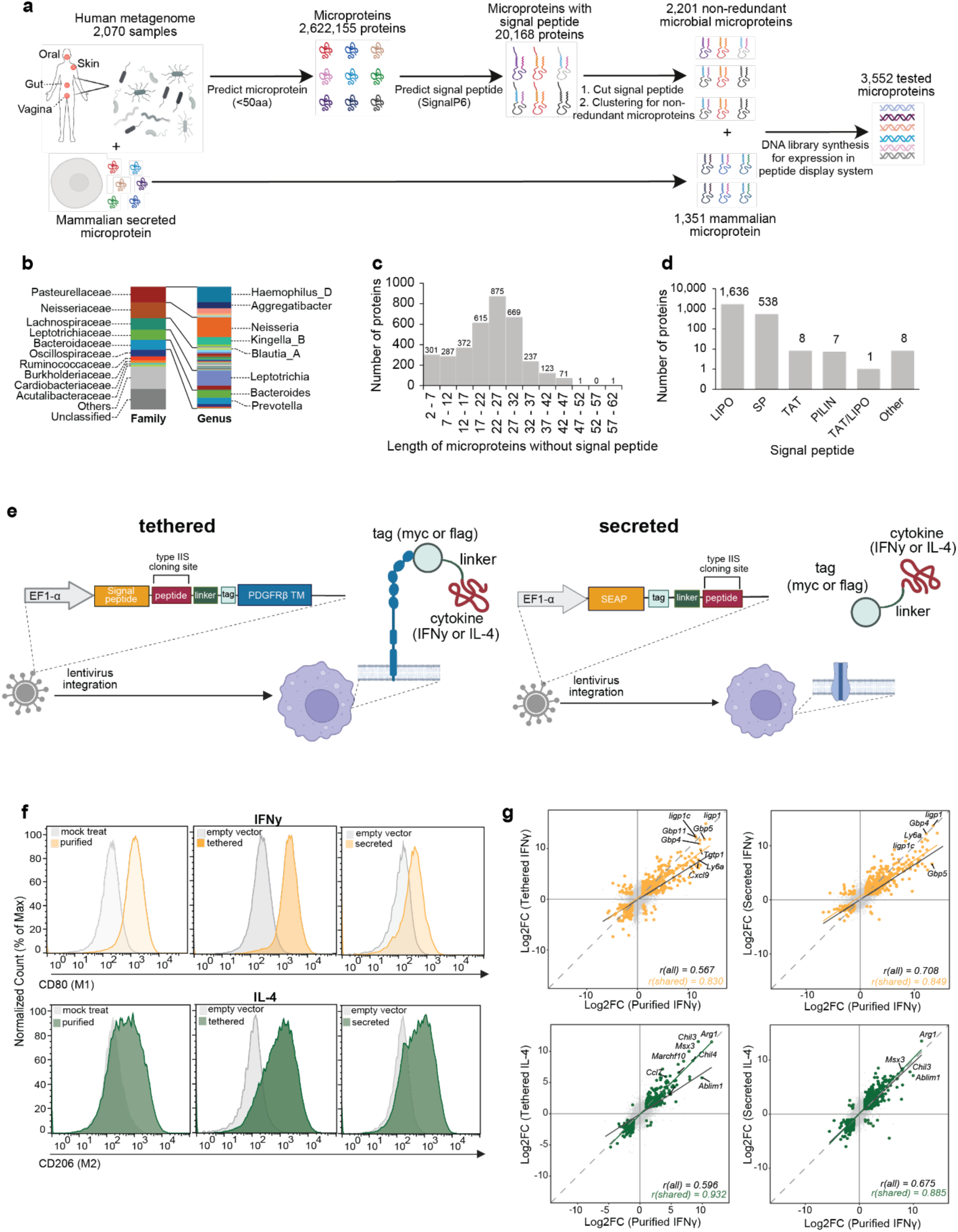
Construction of a microprotein library and recapitulation of macrophage polarization state via cytokine display. a. Workflow for identification of microproteins for further functional characterization b. Distribution at the family and genus level of synthesized microbial microproteins c. Size distribution of synthesized candidate microproteins d. Signal peptide distribution of synthesized microbial microproteins. LIPO= lipoprotein signal peptide. SP= secretory signal peptide. TAT= Twin-Arginine Translocation Signal Peptide. PILIN= Pilin and Pilin-like signal peptide. TAT/LIPO: Twin-Arginine Translocation Lipoprotein Signal Peptide. e. A schematic illustration of the peptide display system. i) For expression of tethered peptides, a vector consisting of an N-terminal signal peptide and peptide insertion cloning site (flanked BsmBI TypeIIS cloning sites) is followed by a flexible linker domain, a myc or flag tag, and a PDGFRB transmembrane domain. ii) For expression of secreted peptides, a vector consisting of an N-terminal secreted alkaline phosphatase peptide (SEAP) signal is followed by an N-terminal myc tag linked to a peptide insertion cloning site (flanked BsmBI sites). Lentiviruses are generated with these engineered constructs and transduced into J774 macrophages. For validation of the display system, either IFN**γ** or IL-4 are cloned into the typeIIS site as cargo and macrophage polarization states were measured. f. Flow cytometry histogram plots of J774 macrophages expressing either tethered or secreted cytokines, as well as purified cytokine controls. Negative controls (grey) compared to IFN**γ** (orange) or IL-4 (green) stimulation, with CD80 and CD206 used as markers of M1 and M2 polarization, respectively g. Pairwise comparison of log2 fold changes for all genes and shared DEGs (IFN**γ**- orange; IL-4; green) in the different stimulation conditions (purified, secreted, and tethered). Linear regression lines for both all genes and for shared DEGs are shown, and Pearson correlation coefficients are indicated.

We find most of the microbial microproteins are derived from either oral or gut associated bacteria, consistent with the input metagenomic distribution (Extended Data Fig. 1a). Taxonomic assignment of these microproteins reveals many are derived from families that colonize the upper respiratory tract and oral cavity (e.g. *Pasteurellaceae*, *Neisseriaceae*, *Leptotrichiaceae*) consistent with the high prevalence of oral commensals in our input data set (Fig. 1b). The size distribution of the microproteins span a range of 2-50 amino acids (except the addition of a 61 amino acid mammalian microprotein: a fragment of Uteroglobulin, a known formyl peptide receptor ligand), with the majority of predicted to be between 17 and 32 amino acids (Fig. 1c). Signal peptide classification predicts that ∼74% of the selected microproteins are surface associated (based on the presence of a lipobox domain), while ∼25% are predicted to be secreted (based on the presence of a standard secretory signal peptide, SP) (Fig. 1d).

### A peptide display system recapitulates macrophage polarization states

To screen thousands of microproteins for immunomodulatory activity, we adapted our recently developed peptide display system^39^ to enable the presentation of microproteins on the surface of macrophages. Briefly, a lentiviral construct encodes a cleavable signal peptide, a peptide cloning site, and a PDGFRβ transmembrane domain to tether peptides at the cell surface (Fig. 1e, tethered); an epitope tag upstream of the transmembrane domain enables surface expression confirmation. To provide a complementary and orthogonal approach for displaying microproteins, we also engineered a platform for peptide secretion, using a secreted alkaline phosphatase peptide signal (SEAP) at the N-terminus connected to a tag and flexible linker (Fig. 1e, secreted).

To assay the functionality of the synthetic tethered and secreted systems in macrophages, we first expressed the well-characterized cytokines IFN**γ** and IL-4, which are known to induce pro- and anti-inflammatory signalling in macrophages, respectively, after exogenous exposure. Upon lentiviral transduction of IFN**γ** and IL-4 expressing constructs into J774 macrophages, we confirmed expression via flow cytometry, with concordance seen between anti-FLAG staining and anti-IL4 staining, respectively (Extended Data Fig. 2a).

As expected, macrophages expressing IFN**γ**, via both the tethered and secreted system, showed increased expression of the pro-inflammatory cell surface markers CD80 and Ly6a by flow cytometry (Fig. 1f, Extended Data Fig. 2b, 2c). Conversely, expression of IL-4 via the display systems leads to increased expression of the anti-inflammatory marker, CD206 (Fig. 1f). To further characterize changes in macrophage cell state, we performed bulk RNA-seq of macrophages expressing IFN**γ** or IL-4 via each display system. Differential gene expression analysis revealed transcriptional signatures consistent with pro- and anti-inflammatory cell states, respectively (Extended Data Fig. 3a, Supplementary Table 2). Principal component analysis (PCA) shows that macrophages expressing cytokines via the tethered and secreted display systems group together with cells exposed to their corresponding purified, exogenously added cytokines, clustering separately from their respective control conditions (Extended Data Fig. 3b). Pearson correlation of log2-fold changes between purified cytokine stimulation and each display modality revealed strong concordance, for both IFN**γ** and IL-4 (Fig. 1g). To identify shared transcriptional pathways, we next performed gene set enrichment analysis (GSEA) across various macrophage perturbation conditions. GSEA revealed consistent enrichment of pro-inflammatory gene sets in macrophages exposed to IFN**γ** across all three delivery modalities, including hallmark IFN**γ** response, TNF-α signaling, and inflammatory response pathways (Extended Data Fig. 3c, Supplementary Table 3). For IL-4, transcriptional changes were similarly concordant, with shared pathway enrichments across delivery modalities (Extended Data Fig. 3c, Supplementary Table 3). Concordance between purified cytokine and the display system for both IFN**γ** and IL-4 demonstrated that the display platform delivers functional cytokine signals capable of altering macrophage transcriptional programs. Lastly, to benchmark our system to physiologically relevant responses, we compared the transcriptional profile of our IFN**γ** exposed macrophages to a curated set of IFN**γ**-responsive genes from stimulated macrophages obtained from *in vivo* perturbational single-cell RNA sequencing (scRNA-seq) data^40^. We found a significant upregulation of these across all three IFN**γ** stimulated conditions (purified, tethered, and secreted) (Extended Data Fig. 3d). Taken together, these findings demonstrate that the peptide display system robustly recapitulates immune modulation in macrophages in a similar manner to exogenous cytokine stimulation.

### A functional genomic screen in macrophages identifies a cluster of pro-inflammatory microproteins encoded in *Leptotrichia*

We next used the tethered peptide system in macrophages to identify candidate immunomodulatory microproteins. We first designed a microprotein library of 7,187 elements (3,552 microproteins, the majority with two coding variants with the exception of some smaller microproteins, as well as 250 random negative control sequences). We cloned this library into the tethered peptide display lentiviral construct and transduced it into J774 macrophages at a low multiplicity of infection (MOI ∼0.2-0.3). To increase our yield of candidate immunomodulatory microproteins, we performed three independent FACS-based screens (each in duplicate) using two independent cell surface markers: CD80, a canonical M1 polarization marker^41^, and Ly6a, a marker also associated with macrophage M1 polarization^42,43^ and one of our most upregulated surface proteins upon IFN**γ** stimulation. To further enhance the sensitivity for identifying immunomodulatory microproteins, the CD80 screen was performed both with and without low dose LPS pre-stimulation, to capture microproteins that could have synergistic effects in the presence of a known pro-inflammatory ligand. For all screens, the top and bottom 10% of cells were sorted; genomic extraction, sequencing and analysis were subsequently performed using the previously developed HT-recruit pipeline to calculate enrichment in the high vs. low populations^44^ (Fig. 2a, Extended Data Fig. 4a).

**Figure 2.**
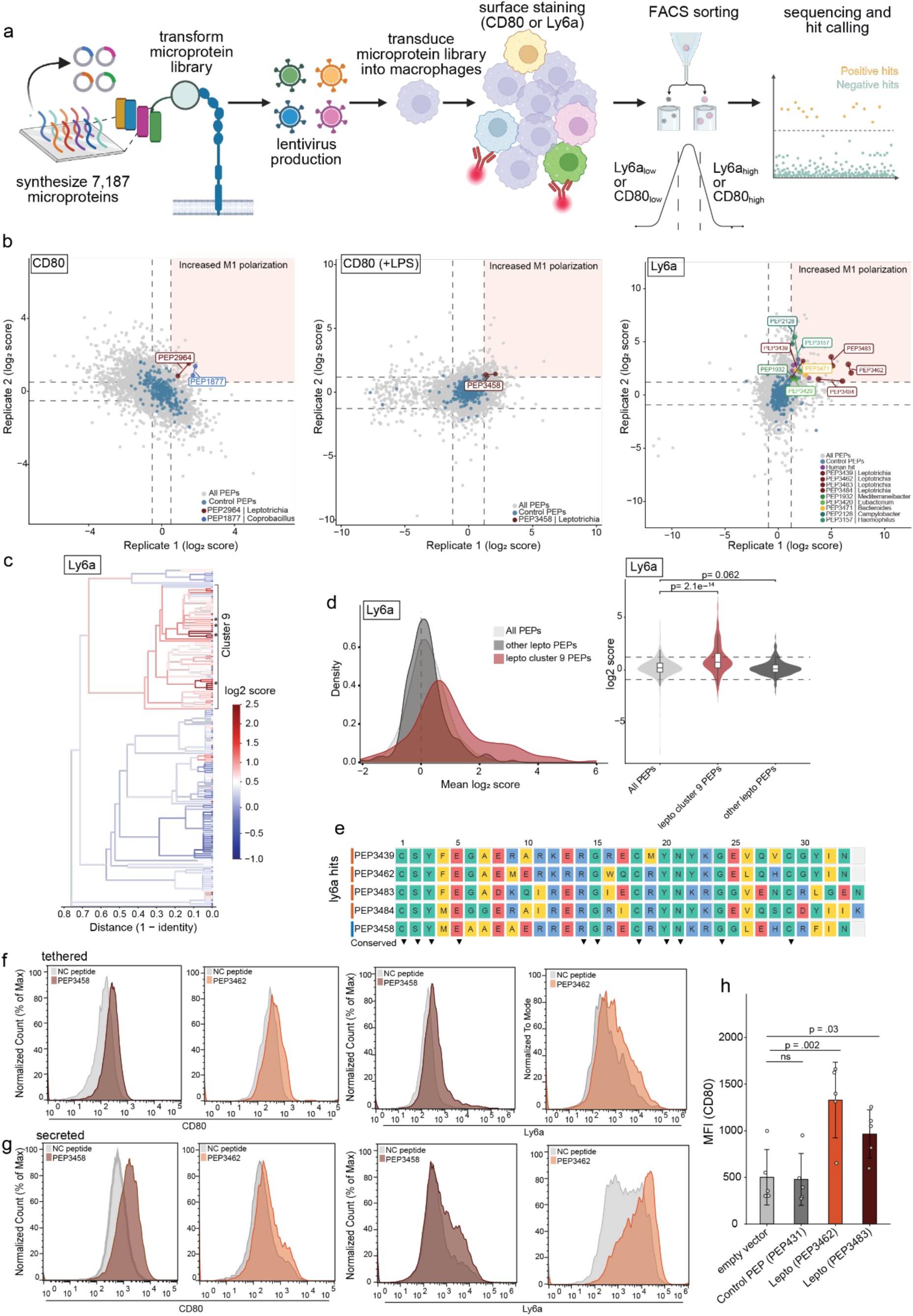
A high-throughput peptide display screen reveals multiple pro-inflammatory microproteins from Leptotrichia. a. Overview of screen design. J774 macrophages were transduced with a library containing microproteins via lentivirus at a low MOI, with greater >1000x library coverage. Cells were then expanded to maintain minimum coverage of library upon cell sorting. The library was then stained with the either CD80 or Ly6a antibody and sorted via FACS. Genomic extraction, library amplification, sequencing and hit-calling were subsequently performed. b. Scatter plot of log₂ enrichment scores for all microproteins screened, from each of the separate sorting screens (i. CD80 staining in absence of LPS, ii. CD80 in presence of LPS, iii. Ly6a). Each axis represents a biological replicate. Values for each element are calculated as log₂ ratio of normalized high counts to low counts (see methods). Dashed lines indicate ±2 standard deviations from the mean with log₂ enrichment scores of random negative control peptides (shown in blue), used as the threshold for hit calling. Colored points are candidate pro- inflammatory microproteins. Candidate hits contain both coding variants and both biological replicates with two standard deviations away from the mean of negative controls. c. Sequence-based clustering of 175 Leptotrichia peptides colored by the Ly6a macrophage polarization screen scores. Hierarchical clustering was performed for all peptide sequences using single-linkage (nearest-neighbor) agglomerative clustering on a pairwise sequence identity distance matrix. Sequence identity between each pair of peptides was computed using a sliding-window best-match approach. Dendrogram branches are colored by the mean Ly6a log₂ score of all leaf nodes below that branch, and each leaf dot is colored by the mean Ly6a score for both biological replicates. Asterisks represent hits from the screen. d. Density plots (left) and violin plots (right) of the mean log₂ enrichment scores (average of replicate 1 and replicate 2) for three microprotein populations: all screened microprotein (excluding *leptotrichia spp*; light grey), Leptotrichia cluster 9 microproteins (red), and all other Leptotrichia microproteins; (dark grey). The violin plot displays the full distribution, with an overlaid boxplot showing the median and interquartile range. Dashed lines indicate ±2 standard deviations from the mean of random negative controls. Statistical comparisons were performed between each Leptotrichia group and the all peptides background using a Mann-Whitney U test. Exact p-values are shown on each comparison bracket. e. Multiple sequence alignment of Leptotrichia clade 9 peptide screen hits. The five Leptotrichia peptides identified as significant hits from the screens (Ly6a- PEP3439, PEP3462, PEP3483, PEP3484; orange) and CD80+LPS (PEP3458; blue). Residues colored by amino acid class: yellow = hydrophobic, teal = polar/small, blue = positively charged, red = negatively charged. Triangles below the alignment indicate identical positions across all five sequences. f. Flow cytometry histogram plots demonstrating increase in CD80 and Ly6a expression in J774 macrophages expressing Leptotrichia pro-inflammatory microproteins (PEP3458, PEP3462) via the tethered display system, in comparison to negative control peptide (in grey). g. Flow cytometry histogram plots demonstrating increase in Ly6a and CD80 expression in J774 macrophages expressing Leptotrichia pro-inflammatory microproteins (PEP3458, PEP3462) via the secreted display system, in comparison to negative control peptide (in grey). h. CD80 median fluorescence intensity (MFI) quantification of flow cytometry assays with tethered peptide display system. Bars represent +/- standard deviation. Peptide expressing strains (PEP431, PEP3462, PEP3483) were compared to the empty vector control using Dunnett’s test following one-way ANOVA. n=5 distinct biological replicates per condition. p- values indicated

To assess the quality of each screen, we evaluated replicate concordance of sorted fraction read counts across all three screens. Count concordance was high across all three screens, for both fraction counts (high and low sorting bins), confirming consistent library representation between replicates (Extended Data Fig. 4b). Comparing the distribution of read counts for each of the four sorted fractions indicated that the peptide library was broadly and evenly represented across sorted populations (Extended Data Fig. 4c). We next sought to determine if any microproteins promoted either a viability defect or a pro-growth phenotype in J774 macrophages. When comparing the input microprotein lentiviral library (plasmid pool) counts to the bulk transduced population in J774 macrophages, we found no microproteins where both coding variants were significantly enriched for either a possible pro-growth effect or viability defect (Extended Data Fig. 4d).

To identify candidate microproteins with pro-inflammatory activity, we required microproteins to have both coding variants and both replicates exceed two standard deviations above the mean log2 score of the random negative controls. As expected, the vast majority of peptides show no significant activity, with these peptides clustering tightly around the distribution of the negative control peptides. Using these criteria, we identified a total of 16 candidate microproteins with putative pro-inflammatory activity in macrophages (Fig. 2b, Extended Data Fig. 5a-c, Table 1, Supplementary Table 4). The majority of microproteins were derived from bacterial species, and most have no known function (Table 1, Supplementary Table 1). One microprotein with increased Ly6a expression, derived from *Bacteroides* species, has sequence homology to the *ompA* family protein (Table 1), a known pathogen-associated molecular pattern (PAMP)^45^.

**Table 1.**
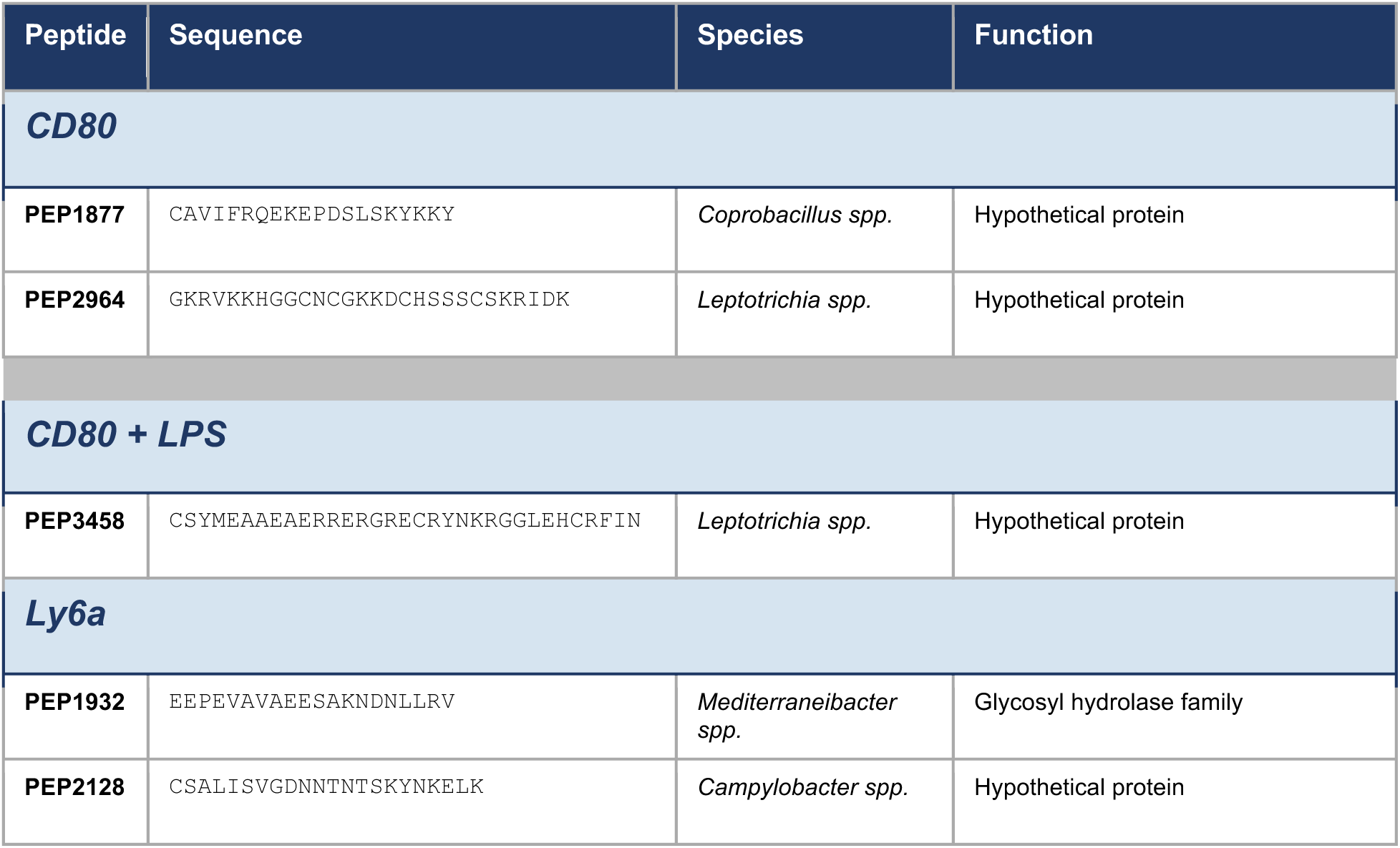
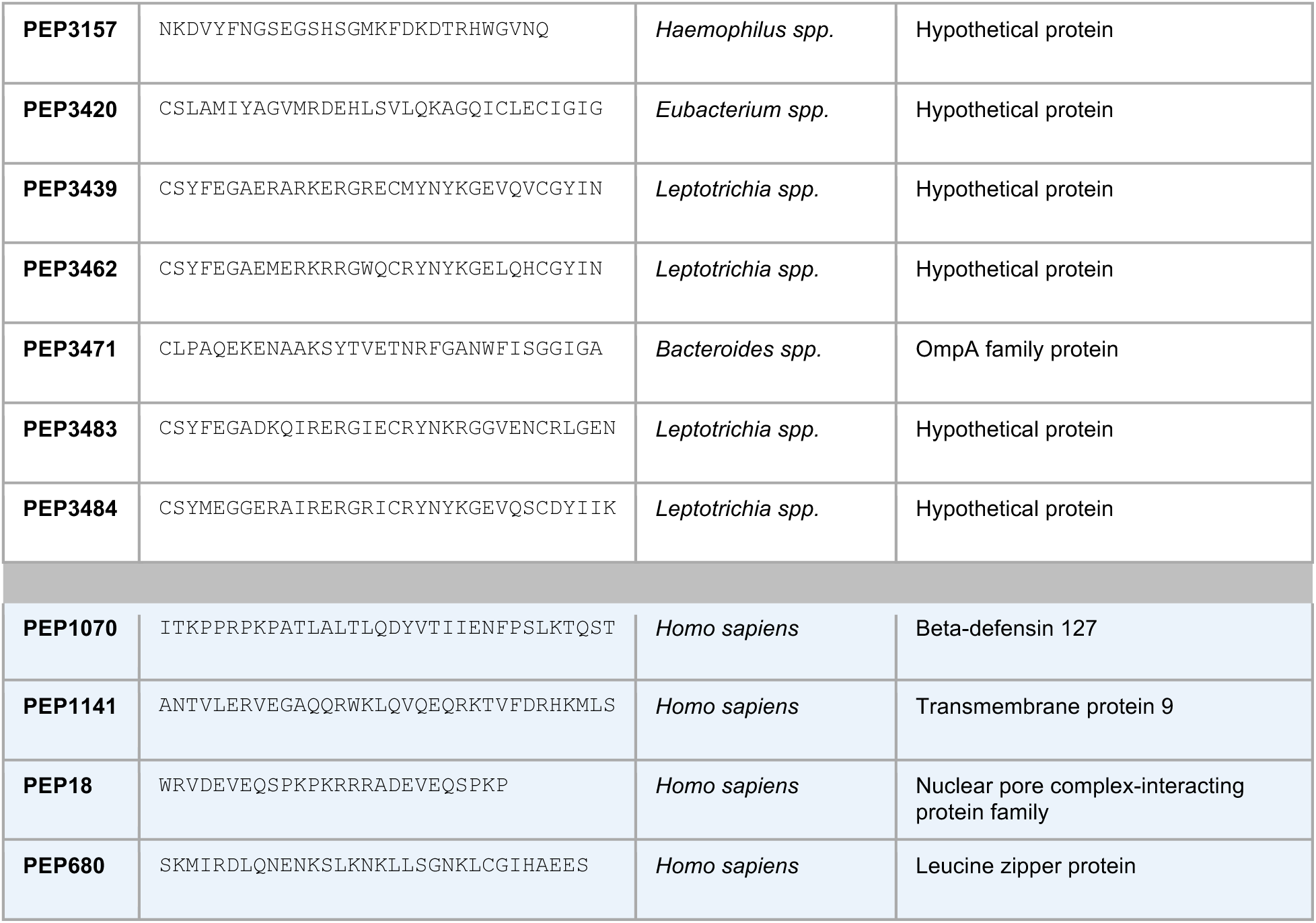
Microbial and human microprotein hits from macrophage polarization screens. Microproteins identified across all three screens (CD80, CD80 with LPS, Ly6a, as indicated). Organism assignment at the genus level for bacterial microproteins. Human microproteins shaded in blue.

Strikingly, across all three screens, we noticed the predominance of *Leptotrichia* microproteins with similar amino acid sequences, which appeared to confer an increase in macrophage polarization, prompting us to further evaluate this observation. To better understand the prevalence of *Leptotrichia* microproteins as candidate hits, we first sought to systematically assess family-level enrichment by calculating the ratio of outlier microproteins for each bacterial family. We found that the Leptotrichiaceae (a member of the phylum Fusobacteriota) was the only significantly enriched taxonomic family in the Ly6a screen, with no families enriched in the CD80 screen (Extended Data Fig. 6a). Given the sequence similarity observed amongst many of the *Leptotrichia* microprotein hits, we hypothesized that a specific cluster of *Leptotrichia* microproteins was responsible for driving the enrichment of immunomodulatory microproteins of the Leptotrichiaceae family. Hierarchical agglomerative clustering of the amino acid sequences of all 175 *Leptotrichia* microproteins in the library, using pairwise sequence identity as the distance metric, identified 11 distinct microprotein clusters (Supplementary Table 5). Cluster 9, containing 65 microproteins, was the primary cluster driving the family-level enrichment of inflammatory microproteins within the screen (Fig. 2c, Extended Data Fig. 6b,c). Cluster 9 *Leptotrichia* microproteins showed significantly elevated screen scores relative to other *Leptotrichia* microproteins and to all other microproteins tested (Fig. 2d, Extended Data Fig. 6d, Supplementary Table 5). Sequence comparison of cluster 9 *Leptotrichia* microproteins suggest an overall highly similar predicted secondary structure, with multiple conserved residues (Fig. 2e). These findings suggest that *Leptotrichia* species contain various microproteins with related sequences which share putative pro-inflammatory activity.

We next validated our screen findings by expressing *Leptotrichia* cluster 9 microproteins in J774 macrophages and assessing pro-inflammatory marker expression via flow cytometry. One of the microproteins selected for validation, the *Leptotrichia* microprotein PEP3458, was identified as a hit in the CD80 screen in the presence of LPS, but not initially a statistically significant hit in the CD80 screen without LPS. Because PEP3458 was in the top 1% of candidate microproteins with increased Ly6a expression, we suspected that while PEP3458 was not a hit in the CD80 screen in the absence of LPS, this finding might have been a false negative. We first expressed PEP3458 via the tethered peptide display system in macrophages, and demonstrated increased expression of CD80 in the presence of LPS, relative to both a control microprotein PEP431 (a matched control based on amino acid length and isoelectric point), and an empty vector control (Extended Data Fig. 6e). We next performed surface staining of Ly6a and CD80 in the absence of LPS, and, indeed, found increased expression in PEP3458-expressing macrophages, relative to controls (Fig. 2f). Expression of two additional cluster 9 *Leptotrichia* microproteins that were hits in the Ly6a screen, PEP3462 and PEP3483, showed an increase in both CD80 and Ly6a expression (Fig. 2f, Fig. 2h). Comparable results were obtained by expressing PEP3458 and PEP3462 in the secreted display system, suggesting that membrane tethering is not necessary for activity (Fig. 2g). To further validate our screen findings, we next tested a hit from an unrelated species, *Coprobacillus* (PEP1877), and also found an increase in CD80 expression (Extended data Fig 6f). In summary, across multiple peptide display screens, we identify multiple bacterial microproteins with candidate M1 polarization activity in J774 macrophages that are validated when tested individually. Most notably, our functional genomic screens and subsequent flow cytometry validation suggests *Leptotrichia* species encode for a cluster of microproteins with similar sequences and M1 polarization activity in macrophages.

### Genomic and structural diversification of the PEP3458 microprotein family in *Leptotrichia*

Given that multiple related *Leptotrichia* microproteins drove a pro-inflammatory macrophage phenotype, we next sought to define the full membership of this family across *Leptotrichia* genomes. We predicted small ORFs from 203 high-quality *Leptotrichia* genomes and performed BLASTP searches using PEP3458, PEP3462, and PEP3483 as references, yielding 512 PEP3458-like ORFs (Extended Data Fig. 7a). Multiple sequence alignment of these 512 ORFs revealed five conserved residues: Cys17, Gly31, Cys34, Gly40, and Cys45 in PEP3458 (Extended Data Fig. 7b, red arrows). In the predicted PEP3458 tertiary structure, Cys17 lies just C-terminal to the signal peptide cleavage site, Gly31 and Gly40 are located in loop regions flanking the β-strand, and Cys34 and Cys45 are positioned on the two β-strands facing each other, likely resulting in a disulfide bond. These placements are consistent with a structural role in maintaining the PEP3458 fold (Extended Data Fig. 7b). On this basis, we defined confident PEP3458-like ORFs as those carrying all five conserved residues and a ColabFold-predicted structure similar to PEP3458, retaining 384 ORFs. Using these as references for iterative homology searches, we ultimately identified 771 PEP3458 family ORFs (Extended Data Fig. 7a).

To characterize the structural diversity of PEP3458 family proteins, we predicted their structures using ColabFold after removing the signal peptide, obtaining models with a mean pLDDT of 80 (Extended Data Fig. 7c). Structural clustering with FoldMason^46^ resolved 10 structurally distinct clades (Fig. 3a). The largest clade, clade 10, which is the only clade that is present in our original microprotein library, contains PEP3458, PEP3462, and PEP3483. Each protein consists of one α-helix and a β-sheet formed by two β-strands. Most other clades share this overall topology but differ in the relative orientation of the α-helix and the β-strands, and in β-strand length. By contrast, clades 01, 05, and 08, all of which encode proteins of greater than 50 amino acids, contain three or more β-strands.

**Figure 3.**
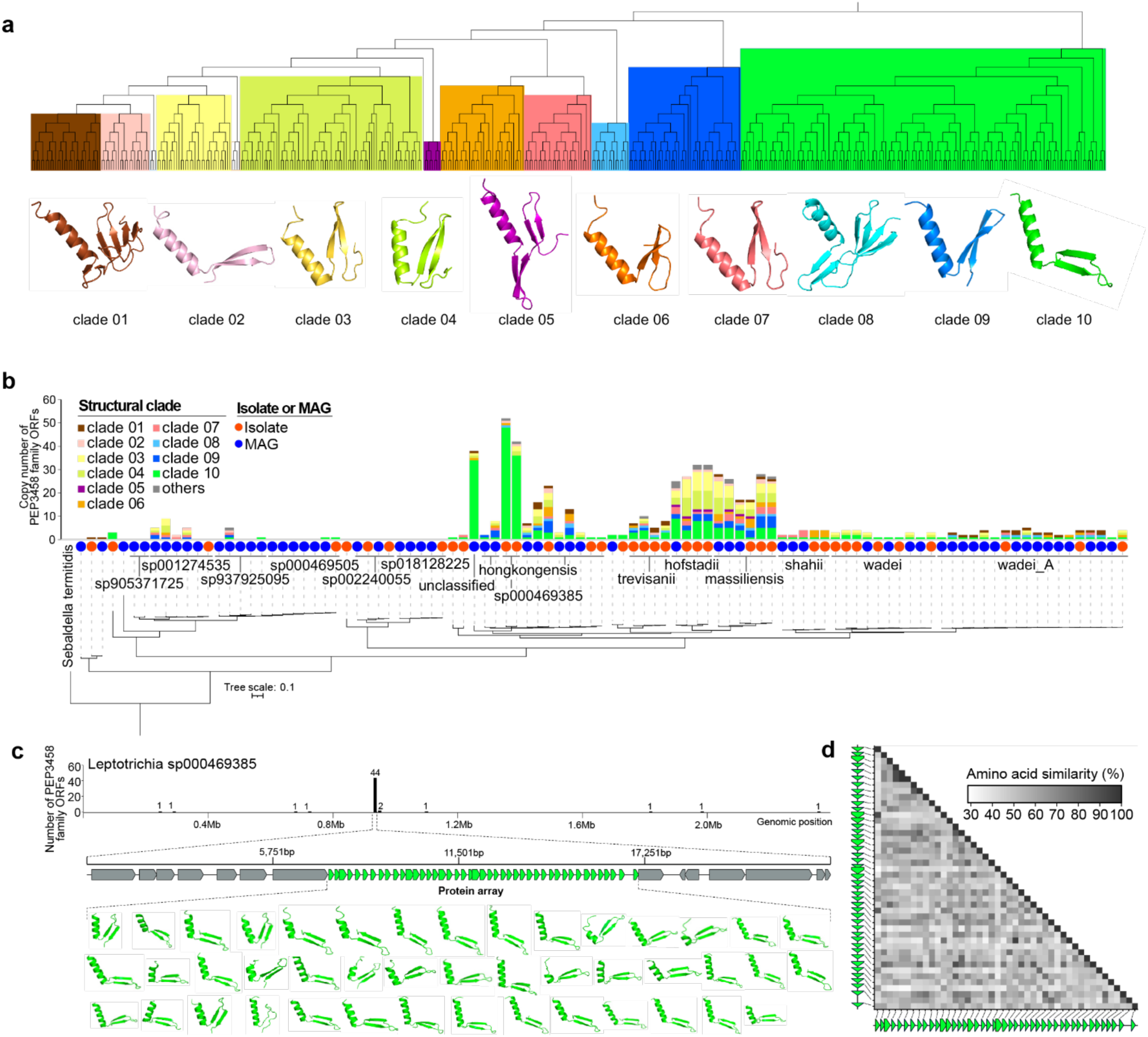
Structural and genetic characterization of PEP3458 family proteins. a. Protein structural diversity of PEP3458 family proteins. Protein structures were predicted with ColabFold and clustered with FoldMason; the dendrogram reflects pairwise structural similarity. Protein structures below the dendrogram show a representative member of each clade, colored as in the dendrogram. b. Copy number of PEP3458 family ORFs in Leptotrichia genomes. The phylogenetic tree was reconstructed using IQ-TREE with 1000 ultrafast bootstrap replicates based on the similarity of core genes obtained using Roary. *Sebaldella termitidis* was used as the outgroup. Red and blue circles represent the origin of genomes from an isolate and MAG, respectively. The GTDB-based species name is shown next to each circle.The bar plot indicates the number of PEP3458 family ORFs in each genome, and the bar color represents the structural clades in the same color as (a). c. Representative genome (*L. sp000469385*) having a large protein array in the genome. The x-axis of the bar plot indicates the genomic position, and the y-axis indicates the number of PEP3458 family ORFs in each 15 kb genomic window. The bottom panel shows a zoom-in of the largest array as a genetic map, with predicted structures of the array proteins below. d. Amino acid similarity within the largest protein array. The heatmap shows the amino acid identity (%) obtained using BLASTP between all combination pairs in the protein array shown in (c).

To investigate the genomic organization of PEP3458 family proteins and the processes that generated their structural diversity, we next quantified the copy number of each structural clade per genome. Several *Leptotrichia* species, most notably *L. massiliensis*, *L. hofstadii*, *L. trevisanii*, *L. hongkongensis*, and *L.* sp000469385, encode a high copy number of PEP3458 family ORFs spanning multiple structural clades (Fig. 3b). In *L.* sp000469385, which carried the highest copy number (53 ORFs), 44 of these ORFs were encoded as tandem repeats in a single genomic locus (Fig. 3c). Hereafter, we refer to a tandem repeat of two or more PEP3458 family ORFs at a single locus as a “microprotein array.” All 44 ORFs in this microprotein array belonged to structural clade 10 (Fig. 3c). Multiple sequence alignment of the 44 ORFs revealed substantial amino acid sequence diversity (Extended Data Fig. 8a). The full length ORFs ranged from 48 to 81 amino acids (Extended Data Fig. 8b), with most of the variation arising from the lipoprotein signal peptide region (Extended Data Fig. 8a). The mean pairwise amino acid identity was 50% (Extended Data Fig. 8c), and pairwise identity showed no clear relationship with the relative position of ORFs within the array (Fig. 3d). In another multi-copy example, the *L. hongkongensis* genome encoded 22 PEP3458 family ORFs distributed as several microprotein arrays of 3–5 ORFs across multiple loci, with each array containing ORFs from distinct structural clades (Extended Data Fig. 8d). Across all 771 PEP3458 family ORFs identified in this study, singletons (single PEP3458 family ORF flanked by non-PEP3458 family ORFs) were the most abundant configuration, accounting for 489 ORFs, whereas the remaining 282 ORFs were encoded within microprotein arrays (Extended Data Fig. 8e). Array sizes varied considerably: two-ORF arrays were the most prevalent (59 arrays), and four arrays contained more than 10 ORFs (Extended Data Fig. 8f).

The above analyses establish that PEP3458 family ORFs are widely distributed across *Leptotrichia* genomes, frequently as microprotein arrays. To determine whether these ORFs are actively transcribed in their native ecological context, we mapped oral metatranscriptomic reads from 19 active root caries samples to the 771 PEP3458 family ORFs. Of these, 78 ORFs received more than 10 mapped reads at >95% sequence identity, indicating detectable transcriptional activity. Notably, two of these transcribed ORFs were co-encoded as a single protein array, with reads mapped to both ORFs and intergenic region at comparable depth, suggesting polycistronic transcription (Extended Data Fig. 8g).

Bacteria frequently encode functionally related genes in close genomic proximity; thus, the functional annotation of genes flanking PEP3458 family ORFs may provide clues to the function of the PEP3458 family itself. We therefore extracted genes flanking PEP3458 family singletons and protein arrays and clustered them by sequence homology. Two clusters were strongly enriched in the vicinity of PEP3458 family ORFs: NADP-dependent glyceraldehyde-3-phosphate dehydrogenase (Cluster_28) and acid phosphatase AphA (Cluster_95) (Extended Data Fig. 9a). By contrast, the largest array (44 ORFs) was flanked by *treC* (α,α-phosphotrehalase) and *cinA* (an ADP-ribose pyrophosphatase involved in DNA damage and competence), and this configuration was conserved across all *Leptotrichia* genomes carrying large protein arrays (Extended Data Fig. 9b). Because *treC* and *cinA* are also adjacent in *Leptotrichia* species that lack PEP3458 family protein arrays (Extended Data Fig. 9c), these results suggest that the *treC– cinA* intergenic region serves as a genomic site for the acquisition and expansion of PEP3458 family protein arrays, either through insertion of pre-assembled array cassettes, gene amplification, or a combination of both.

### *Leptotrichia* microproteins induce a pro-inflammatory state in macrophages

To further delineate the effect of *Leptotrichia* microproteins on macrophage cell state, we performed bulk RNA-seq of macrophages expressing the tethered *Leptotrichia* microprotein PEP3458. We first compared macrophages expressing a control peptide, PEP431, to those expressing the empty vector, and found only one differentially expressed gene (DEG), confirming that the expression construct with the control peptide PEP431 does not significantly alter the transcriptional state. Comparison of PEP3458-expressing macrophages (both with and without low dose LPS) to PEP431-expressing controls, identified over 200 DEGs, including pro-inflammatory cytokines, chemokines, and macrophage activation markers (e.g. *Marco*, *Ccl7*, *Ccl2*) (Figure 4a, Extended Data Fig. 10a, Supplementary Table 6).

**Figure 4.**
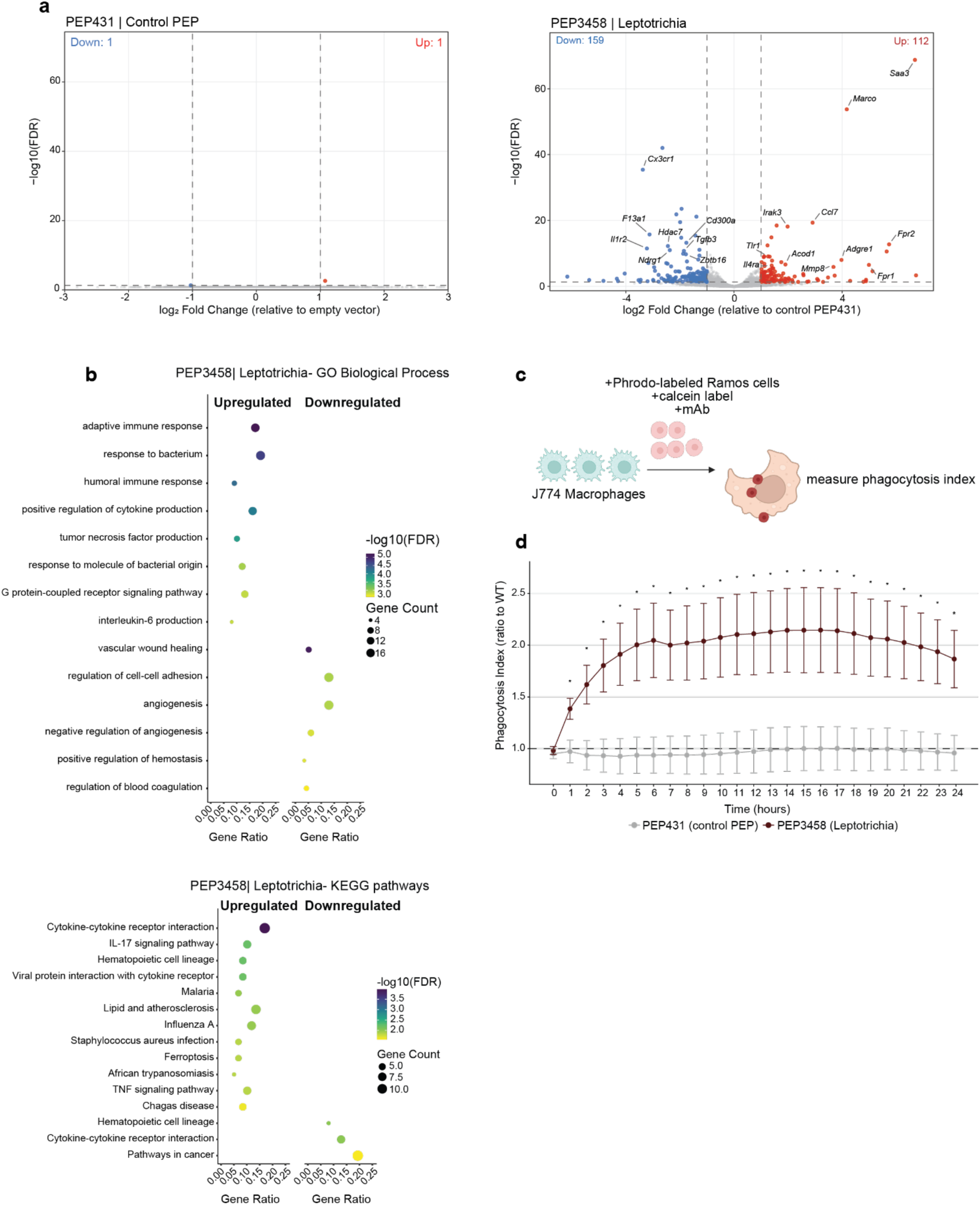
Leptotrichia microproteins induce broad immunomodulatory changes in macrophages. a. Volcano plot of bulk RNA-seq from J774 macrophages expressing the PEP431 control microprotein (left) to empty vector, and the Leptotrichia microprotein, PEP3458, relative to the PEP431 control microprotein (right). n= 3 distinct biological replicates for all conditions. Red dots represent genes differentially upregulated, blue dots represent genes differentially downregulated. Select individual genes associated with M1 polarization labeled. b. (top) GO Biological Process dot plot showing top enriched pathways per direction. Dot size represents gene count; color intensity indicates statistical significance (-log10 FDR). Upregulated pathways include M1-associated processes such as tumor necrosis factor production, NF-κB signaling, and antimicrobial responses. Downregulated pathways include M2- associated wound healing and angiogenesis processes. (bottom) KEGG pathway enrichment showing upregulation of cytokine signaling and IL-17 pathways. Redundant GO terms were filtered using semantic similarity based on GO hierarchical relationships (see methods); complete unfiltered results are provided in Supplementary Table 9. c. Schematic of co-culture of phagocytosis assays. J774 macrophages transduced with peptide display constructs (empty vector, negative control PEP431, or Leptotrichia PEP3458) are co-cultured with pHhrodo red labeled Ramos lymphoma cells. d. Phagocytosis assay quantifying pHrodo red signal over time after co-culture of J774 macrophages expressing with Ramos cells with selected time points for time course. Cells expressing either PEP431 (grey) or PEP3458 (red) are normalized to J774 macrophages expressing empty vector. n= 6 distinct biological replicates for all conditions. One-sample t-test performed for statistical analysis (* p<0.05).

To determine whether microproteins from phylogenetically divergent bacteria induce distinct transcriptional programs, we performed RNA-seq from macrophages expressing a separate candidate immunomodulatory microprotein - PEP1877 from *Coprobacillus*. We identify over 1,000 DEGs in macrophages expressing PEP1877, with PCA demonstrating that macrophages expressing PEP3458 or PEP1877 occupied distinct transcriptional states (Extended Data Fig. 10b,c, Supplementary Table 6).

To define the transcriptional states induced by PEP3458, with and without LPS, we performed over-representation analysis (ORA) of DEGs. Analysis of both Gene Ontology (GO) biological processes and KEGG pathways revealed an upregulation of inflammatory pathways, including TNF signaling, IL-1 response, and IL-17 signaling, and a downregulation of pathways typically associated with M2 polarization, such as wound healing and angiogenesis, suggesting a transcriptional shift away from anti-inflammatory and tissue-repair programs (Fig. 4b, Extended Data Fig. 10d, Supplementary Table 7). ORA of PEP1877 showed a distinct profile of upregulated and downregulated immune pathways (e.g. T cell differentiation, TNF alpha signalling pathway, response to TGFβ), as well as non-immune pathways (circadian rhythm, mineral absorption) (Extended Data Fig. 10e, Supplementary Table 7). The enrichment of non-immune pathways such as circadian rhythm and mineral absorption suggests that *Coprobacillus* PEP1877 may engage broader metabolic or homeostatic programs beyond classical immune activation.

We next sought to assess the functional consequences of *Leptotrichia* microprotein expression in macrophages. Given the pro-inflammatory transcriptional state induced by PEP3458, we hypothesized that PEP3458-expressing macrophages would exhibit enhanced phagocytic capacity. To test this, we co-cultured J774 macrophages expressing PEP3458 with Ramos B lymphoma cells and measured phagocytic uptake of the Ramos B lymphoma cells (Fig. 4c). We found that macrophages expressing PEP3458 exhibited increased phagocytosis relative to the control peptide, PEP431 (Fig. 4d). Together, these findings demonstrate that *Leptotrichia* microproteins induce a pro-inflammatory transcriptional program in macrophages with functional consequences for phagocytic activity, and that microproteins from distinct bacterial species drive divergent macrophage responses.

### *E. coli* expressing *Leptotrichia* microproteins promote an inflammatory response in macrophages

We next sought to test the response of *Leptotrichia* PEP3458-family microproteins in the context of a bacterial co-culture model where a bacterium expressing the microproteins in ‘trans’ was exposed to J774 macrophages. We used the endotoxin-free ClearColi expression system^47^, to minimize the significant inflammatory response generated by the presence of LPS-containing *E. coli*. ClearColi strains were transformed with a medium-copy plasmid (pCDF) containing microproteins expressed under a medium-strength constitutive promoter (J23105) (Fig. 5a), either with or without a C-terminal 6x his tag, and co-cultured with J774 macrophages for subsequent marker expression and transcriptomic profiling. Western blot analysis demonstrated no clear differences in expression level across all microprotein expressing strains (Extended Data Fig. 11a). We next tested the macrophage response to ClearColi strains expressing two *Leptotrichia* PEP3458-family microproteins, PEP3462 and PEP3483 (with the exclusion of PEP3458 due to poor expression via western blot), and compared it to both the empty vector strains as well as a length- and charge-matched negative control bacterial microprotein (PEP3473) with no immunomodulatory activity in the performed screens. Flow cytometry of macrophages exposed to ClearColi strains expressing *Leptotrichia* microproteins PEP3462 or PEP3483 showed increased expression of CD80 relative to both empty vector and PEP3473 for both His and non-His tagged expressing strains, confirming the epitope tag does not influence the observed response, with isotype control showing no relative difference (Fig. 5b, Extended Data Fig 11b-d). Bacterial CFU counts at the point of infection and at the time of harvest did not differ significantly between strains, indicating that differential CD80 expression was not driven by variation in bacterial replication (Extended Data Fig. 11e).

**Figure 5.**
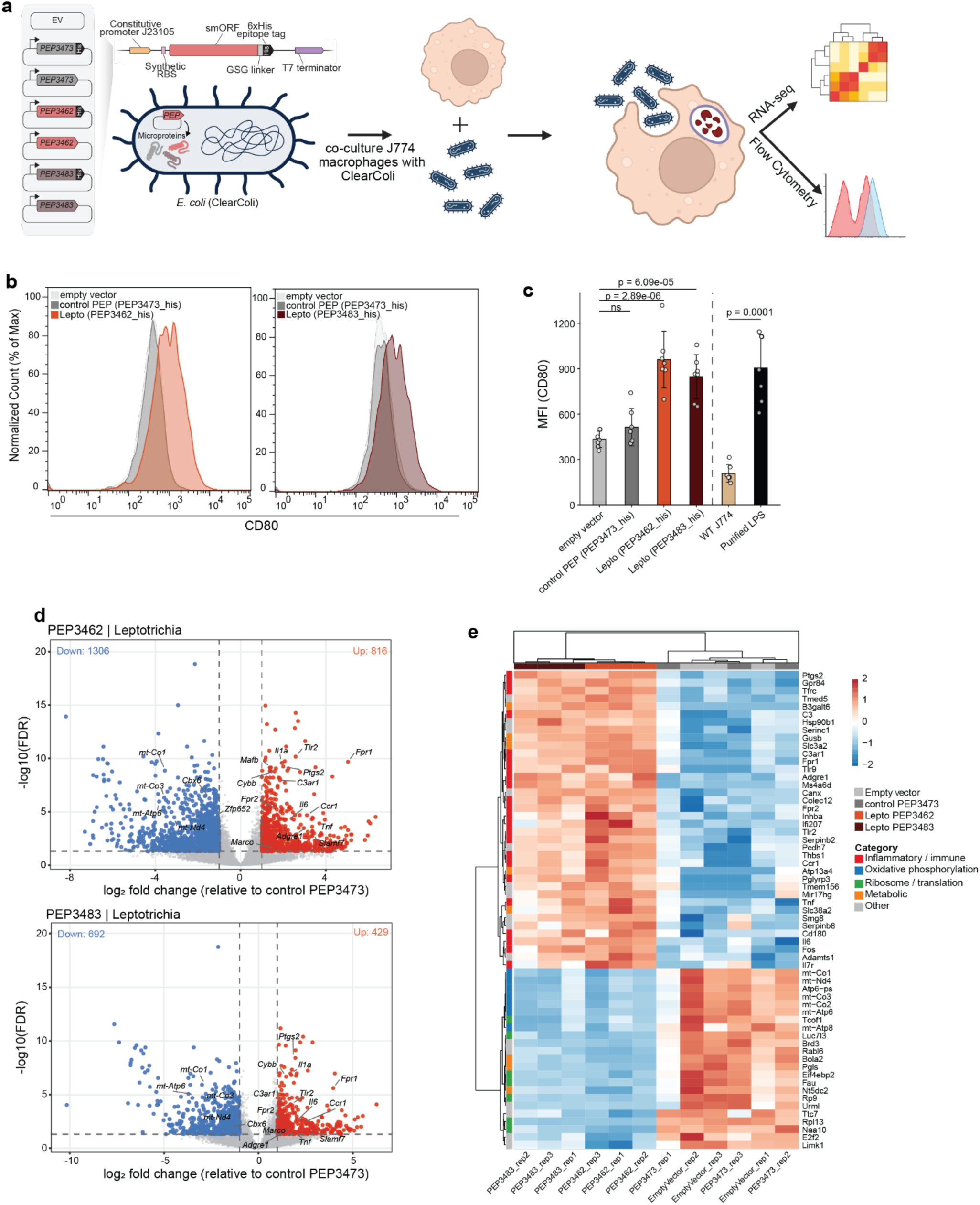
Macrophages exposed to ClearColi expressing Leptothrichia microproteins exhibit increased inflammatory response. a. Schematic of co-culture of J774 macrophages with ClearColi. ClearColi strains were transformed with expression vectors encoding Leptotrichia microproteins (PEP3462 or PEP3483), and compared to a control peptide (PEP3473). ClearColi strains were incubated with macrophages and harvested for flow cytometry and transcriptomics analyses. b. Flow cytometry histograms demonstrating increased CD80 expression for macrophages exposed to ClearColi strains expressing Leptotrichia microproteins PEP3462 (left, orange) and PEP3483 (right, red), relative to empty vector and negative control peptide (grey), PEP3473. c. CD80 median fluorescence intensity (MFI) quantification of flow cytometry assays from ClearColi macrophage co-culture assays. Bars represent +/- standard deviation. Peptide expressing strains (PEP3462, PEP3483, PEP3473) were compared to the empty vector control using Dunnett’s test following one-way ANOVA. WT J774 and purified LPS were compared by Welch’s unpaired t-test. n=6 distinct biological replicates per condition. p-values indicated. d. Volcano plot of bulk RNA-seq from J774 macrophages co-cultured with ClearColi strains. Comparison of Leptotrichia microproteins PEP3462 (left) and PEP3483 (right), relative to control microprotein PEP3473. n=3 distinct biological replicates per condition. Red dots represent genes differentially upregulated, blue dots represent genes differentially downregulated. e. Heatmap showing DEGs in J774 macrophages co-cultured with ClearColi expressing PEP3462 or PEP3483 relative to the negative control peptide, PEP3473. Gene selection criteria: upregulated genes padj < 0.001, |log2FC| > 1.5; downregulated genes padj < 1×10⁻⁵, |log2FC| > 1.5. Each row represents an individual gene, z- scored across samples. Column colors indicate experimental condition: light grey, Empty Vector; dark grey, PEP3473 (negative control peptide); orange, PEP3462; dark red, PEP3483. Row annotation bar indicates functional gene category.

To further delineate the response of macrophages to ClearColi expressing *Leptotrichia* microproteins, we performed RNA-seq of macrophages co-cultured with strains expressing PEP3462 or PEP3483. As expected, exposure of J774 cells to ClearColi induced upregulation of genes associated with M1 polarization, with upregulation of pathways involved in bacterial infection and downregulation of pathways involved in chromosome segregation and DNA replication (Extended Data Fig. 12a-b, Supplementary Table 8). We observe minimal DEGs between exposure to ClearColi expressing empty vector and the PEP3473 control microprotein, confirming no increased immunomodulatory activity from non-specific microprotein expression (Extended Data Fig. 12c, Supplementary Table 8). Transcriptomic analysis of macrophages co-cultured with ClearColi that are expressing the *Leptotrichia* microproteins PEP3462 and PEP3483 showed a significantly increased expression of a suite of M1-associated inflammatory genes (*Fpr1, Fpr2, MARCO, TLR2)* and decreased expression of transcriptional regulators associated with anti-inflammatory macrophage states, including *Nfix* and *RelB,* an NF-κB family member that epigenetically silences pro-inflammatory gene promoters during inflammatory resolution^48–50^. (Fig. 5c, Extended Data Fig. 12d, Supplementary Table 8). Notably, cells co-cultured with PEP3462- or PEP3483-expressing ClearColi also showed decreased expression of mitochondrial oxidative phosphorylation genes (*mt-Co1*, *mt-Co2*, *mt-Nd4*, *mt-Atp6*), consistent with the metabolic reprogramming characteristic of committed M1 macrophage polarization (Fig. 5c, Supplementary Table 8)^51^. ORA corroborated these findings, identifying enrichment of inflammatory response and innate immune signaling pathways among upregulated DEGs, and depletion of metabolic and homeostatic pathways among downregulated genes (Extended Data Fig. 12e, Supplementary Table 9). Together, these data demonstrate that *Leptotrichia* microproteins drive a robust M1-like transcriptional program in macrophages, encompassing both pro-inflammatory gene activation and metabolic reprogramming, when delivered in the physiologically relevant context of bacterial infection.

## Discussion

The systematic annotation of microproteins across genomes has revealed a vast and largely uncharacterized layer of the proteome^25,26,52^. However, the experimental tools required to systematically screen bacterial microproteins for host-modulatory activity have lagged behind the pace of computational discoveries. In this work, we expand the genomic toolkit of microprotein discovery by developing and validating a peptide display platform capable of expressing microproteins at the macrophage surface, using it to functionally screen microbial small proteins for cross-kingdom immunomodulatory activity.

The peptide display platform described here provides a scalable cell-based strategy for the systematic functional annotation of bacterially encoded proteins. The pooled format of this screen enables simultaneous evaluation of thousands of candidate bacterial sequences, providing a throughput advantage over smaller scale approaches to annotate uncharacterized microproteins, which is particularly appealing given the scale of metagenomic datasets^35^. This modular system is, in principle, adaptable to diverse cell types, allowing for discovery of microbial factors that could, for example, alter epithelial cell function or lymphocyte activation. While the display system is currently limited to genetically encoded protein sequences and does not capture immune-active bacterial lipids or carbohydrates, recent technical advances enabling the affordable synthesis of longer sequences expands the screenable sequence space to include protein domains and full-length proteins beyond the microprotein range. The display platform should also be compatible with diverse functional readouts, including flow cytometry-based screens for surface marker expression, functional assays such as phagocytosis or cytokine secretion, and more unbiased approaches such as single-cell RNA sequencing. Lastly, the continued expansion of metagenomic databases considerably increases the pool of candidate proteins, both small and large, available for functional annotation. The identification of small proteins with immunomodulatory activity could serve as a platform for the development of peptide-based therapeutics that target innate immune pathways^53^.

Upon screening thousands of microproteins for immunomodulatory activity, we identified a unique cluster of related small proteins encoded by *Leptotrichia* species, Gram-negative anaerobic commensals that colonize the human oral cavity. *Leptotrichia* are members of the Fusobacteriales order, which includes the oncogenic bacterium *Fusobacterium nucleatum*; despite this association, little is known about how *Leptotrichia* colonization impacts human health. As known members of the healthy oral microbiome, metagenomic studies demonstrate that *Leptotrichia* are notably enriched in subgingival niches^54,55^. The close contact with gingival epithelium and tissue-resident macrophages may position them to engage host innate immune pathways through direct cell-cell contact or secreted factors. Intratumoral *Leptotrichia* abundance has been identified as a favorable prognostic marker in head and neck squamous cell carcinoma (HNSCC), with higher *Leptotrichia* levels associated with earlier tumor staging and improved clinical outcomes, leading to the hypothesis that *Leptotrichia* could exert anti-tumor activity via stimulating or specifically modulating host immune responses^56,57^. Beyond HNSCC, *Leptotrichia* has been detected within the tumor microenvironment of colorectal cancer, where its presence has been linked to alterations in macrophage state, further supporting a broader role for this genus in shaping innate immune responses at mucosal sites^58^.

Although the chromosomal location of microprotein arrays varies across *Leptotrichia* genomes, we consistently observe that the longest arrays are encoded between predicted *treC* and *cinA* genes, suggesting a possible link between microprotein expression and trehalose metabolism and/or the bacterial stress response. Trehalose lipids are found in the cell walls of Gram-positive and Gram-negative bacteria, and are known modulators of host immunity^59^. Notably, the glycolipid trehalose 6,6 dimycolate (TDM) protects *Mycobacterium tuberculosis* from macrophage killing via inhibition of phagolysosomal fusion^60^. TDM activates macrophages with expression of inflammatory cytokines^61^, with proposed binding to C-type lectin receptors Mincle^62^ and Dectin 3^63^, as well as the MARCO receptor^64^. Related trehalose phospholipids have been reported in other bacterial pathogens and similarly demonstrate the capacity to activate the Mincle receptor^65^. Furthermore, trehalose accumulation and the metabolic machinery involved in its processing have been implicated in virulence, host colonization, and stress tolerance across diverse bacterial species^59^. Consequently, understanding the regulation of PEP3458-family microprotein expression in a native ecological context, and how microprotein expression influences the immunological and clinical phenotypes observed in human cancers, remains an intriguing avenue for future investigation.

The finding that *Leptotrichia* encodes for extended arrays of structurally related sequences provides important context for the evolutionary origin of these microproteins. We hypothesize that PEP3458-family microprotein arrays evolved through gene amplification and divergence, a known mechanism across diverse species and contexts^66^. Amplification occurs through a range of proposed molecular mechanisms, including RecA-dependent recombination between repeated sequences and replication-based processes that operate without requiring sequence homology at duplication junctions^67^. Interestingly, we note that overall array members do not share a consistent chromosomal location, and that we do not observe direct repeats flanking array members. Overall, the genomic organization of these arrays is therefore most consistent with mechanisms that do not leave classical recombination signatures, though the available data do not allow us to resolve the specific mechanism of expansion. Regardless, the retention of conserved structural motifs across array members despite substantial primary amino acid sequence divergence suggests that functional properties have been preserved throughout their evolutionary history, a pattern consistent with selection acting to retain the shared structural scaffold. The combination of sequence diversity and structural conservation distinguishes these arrays from alternative scenarios such as recent duplications, which would retain high sequence similarity, and copies that diverged without functional constraint, which would be expected to accumulate loss-of-function mutations and gradually lose function through mutational decay.

Overall, we hypothesize that the selective pressures driving microprotein array expansion in *Leptotrichia* arise from ongoing evolutionary conflict between commensal bacteria and host immune systems, a dynamic consistent with Red Queen coevolution^68^. Expansion and diversification of host-interaction gene families under immune-related selection is a recurring genomic response across diverse organisms. In *Staphylococcus aureus*, tandem gene amplifications arise under host immune and antimicrobial selection, and in vaccinia virus, the antihost gene K3L undergoes copy number expansion under experimental selection in human cells, facilitating the acquisition of adaptive mutations that defeat host antiviral defenses ^69,70^. Such gene accordions may be a critical and underappreciated feature of many genetic conflicts involving strong selection, in which transient amplification events are difficult to detect, yet play a critical role in driving adaptation. We propose that the PEP3458-family microprotein array represents a response to genomic conflicts. A structurally conserved but sequence-diverse repertoire of microproteins may collectively broaden *Leptotrichia’s* capacity to engage immune pathways within the heterogeneous landscape of the oral mucosa. The functional scope of individual array members, the degree to which they act in concert, and the broader biological roles these microproteins may play - including potential interactions with competing members of the oral microbiome - remain compelling questions that future comparative genomic and functional studies will be well-positioned to address.

This study has several limitations. First, constraints of the design strategy mean the display system may lead to false negatives in a pooled screening format. In the current construct design, the N-terminus is constrained by its attachment to the signal peptide and tethering domain, which may limit the activity of microproteins that require a free N-terminus for receptor engagement or signaling. Future studies using pooled varying vector designs (e.g. the secreted display system) could enhance the sensitivity of bioactive microproteins. Second, expression of bacterial microproteins in eukaryotic cells limits our ability to account for naturally occurring post-translational modifications such as glycosylation, methylation, cyclization or other specific processing, which in many cases are critical for the activity of bacterial membrane-associated proteins. Finally, while the synthetic display approach enables the discovery of putative immunomodulatory sequences, experimental approaches to assess native microprotein function within the bacterium remain challenging for organisms that are difficult to culture or genetically manipulate.

This work establishes that bacterially encoded microproteins can directly reprogram immune cell states, and provides a scalable platform for their systematic discovery. Our findings emphasize the unappreciated repertoire of microbiome factors with potential relevance to cancer, infection, and inflammatory disease. Continued functional annotation of the microbial microproteome, guided by the genomic and experimental framework developed here, could ultimately reveal new targets for microbiome-based therapeutic intervention.

## Methods

### Identification of the comprehensive secreted or membrane-associated microproteins

Publicly available microprotein sequences derived from the Human Microbiome Project were obtained from the supplementary table of Sberro *et al*^25^. Two additional human metagenomic short read sequence datasets (Gopalakrishnan *et al* and Franzosa *et al*) were used for additional identification of the microproteins^16,36^. Human microproteins were directly obtained from Foster *et al*.^37^ with an additional list of manually curated microproteins incorporated based on proposed GPCR binding and/or immunomodulatory activity (see Supplementary Table 1 for details). For bacterial microproteins, reads were subjected to quality filtering (https://github.com/bhattlab/bhattlab_workflows) as follows: briefly, reads were deduplicated using HTStream SuperDeduper (v.1.3.3), and low-quality bases were trimmed using TrimGalore (v.0.6.7). The quality-filtered reads were mapped to the human genome (hg19) using bwa^71^ (v.0.7.17 31), and all mapped reads were removed. Then, the short reads were assembled using MEGAHIT (v1.2.9)^72^. Microprotein sequences were predicted using smORFinder (v1.0.0)^73^, and deduplicated using mmseq2 (v13.45111)^74^ with 100% amino acid identity over 100% of the sequence length. Thereafter, this collection of microproteins was subjected to SignalP6^38^ to detect predicted signal peptides; the portion of the sequence C-terminal to the predicted transmembrane or secretion tag site was retained. Finally, these microprotein sequences were again deduplicated using mmseq2 (v13.45111) with 100% amino acid identity. A subset of peptides obtained from Sberro et al. contained one or more ‘X’ residues, which Prodigal assigned to codons containing ambiguous bases during translation. To produce a fully defined amino acid sequence set for the final library, all ‘X’ residues were substituted with glycine (G).

### Taxonomic annotation of microproteins

All microproteins were aligned to the amino acid sequences of the Progenome3 database^75^ with taxonomic annotation based on GTDBtk (v2.5.2)^76^ using BLASTP (v2.2.31+)^77^ with >50% identity and >50% query coverage. The best bit score alignment was identified as a potential origin genome of each microprotein. When multiple alignments shared the highest bit score across different genomes, the lowest common ancestor across those matched genomes was determined.

### Identification of the PEP3458 family proteins from Leptotrichia genomes

All *Leptotrichia* genomes in the NCBI database (as of 02/06/2025) were downloaded. The genome contamination rate was assessed using CheckM2 (v1.0.2)^78^ and genomes with >5% contamination were discarded. Additional high-quality *Leptotrichia* genomes were also collected from metagenomic datasets, as follows: long read metagenomic sequences from a salivary microbiome dataset^79^ were assembled using metaFlye (v2.9.2-b1786)^80^ using default parameters. The resultant contigs were binned using SemiBin2 (v2.0.2)^81^ with single sample binning mode and human_oral option. *Leptotrichia* bins were identified using GTDBtk (v2.4.0). The completeness and contamination of the *Leptotrichia* chromosomal bins were determined using CheckM2 (v1.0.2) and bins with >90% completeness and <5% contamination were subjected to further analysis. Open reading frames (ORFs) were predicted using Prodigal (v2.6.3)^82^ with meta option. ORFs with <100 amino acids were aligned to PEP3458, PEP3462, and PEP3483 sequences using BLASTP (v2.2.31+). Initially, all proteins which satisfied >20% amino acid identity and >50% query coverage were identified as PEP3458-like proteins. Multiple sequence alignment was obtained from all candidates of the PEP3458-like proteins and PEP3458, PEP3462, and PEP3483 sequences using MUSCLE (v3.8.31)^83^. From the multiple sequence alignment, all proteins having the five amino acid residues that were conserved among this family (Cys17, Gly31, Cys34, Gly40, and Cys45 in PEP3458, PEP3462, and PEP3483) were selected as ‘confident’ PEP3458 family proteins. Then, to maximize the number of PEP3458 family proteins, all ORFs with <100 amino acids were aligned to confident PEP3458 family proteins using BLASTP (v2.2.31+) with >50% identity and >90% query coverage. Newly identified hits were added to the reference set, and the search was repeated until no further hits were recovered. Finally, protein structures of all small proteins aligned with PEP3458 family proteins were predicted using ColabFold (v1.5.2)^84^ with --msa-mode mmseqs2_uniref_env --model-type auto --num-models 1. Proteins with structures different from PEP3458 were manually discarded based on visual inspection.

### Structural clustering of the PEP3458 family proteins

Signal peptides of all PEP3458 family proteins were predicted using SignalP5, which in our analysis had a lower false positive rate than SignalP6, which was used in the initial microprotein library development. As before, the portion of the sequence C-terminal to the predicted transmembrane or secretion tag site was retained. All resultant PEP3458 family proteins were clustered with 100% identity and 100% identical sequence length using CD-HIT (v4.8.1)^85^ to generate a nonredundant set. Protein structure was predicted using ColabFold (v1.5.2) with -- msa-mode mmseqs2_uniref_env --model-type auto --num-models 1. Then, pdb files were used for creating the protein structure-based phylogenetic tree using FoldMason^86^ (https://search.foldseek.com/foldmason) using default settings.

### Genomic analysis of Leptotrichia

The genome of the isolated *Leptotrichia* (GCA_000469385.1) was assembled using Flye (v2.9.2-b1786). The *Leptotrichia* microprotein array was discovered through manual searches of the gene coordinates of the PEP3458 family microproteins within each *Leptotrichia* genome and metagenome assembled genome. Genomic visualization of the protein arrays was performed using LoVis4u (v0.1.5)^87^. Amino acid similarity within the protein array was determined using BLASTP (v2.2.31+). For the phylogenetic tree of the *Leptotrichia* genomes, multiple sequence alignment of a concatenated list of core genes was obtained using roary (3.13.0)^88^ with -i 90 option, and the phylogenetic tree was constructed using IQ-TREE (3.1.1)^89^ with the TIM2+F+I+R4 model that was determined by ModelFinder Plus in IQ-TREE. The bootstrap value was obtained by IQ-TREE with -B 1000 option. For the identification of genes that are enriched in the locations up and downstream of the PEP3458 family ORFs, ORFs located one position upstream and downstream of each PEP3458 family ORF and protein array were extracted. The extracted neighborhood genes were clustered using CD-HIT(v4.8.1) with -c 0.9 - n 5 -aS 0.5 -aL 0.5 options. Functional annotation of these neighborhood genes was performed using eggNOGmapper (v2.1.6)^90^. Synteny plots for vicinity regions of PEP3458 family ORFs were obtained using lovis4u (v0.1.5) and AliTV^91^.

### Metatranscriptome analysis of protein arrays

Metatranscriptomic sequence data from active root caries was downloaded (Accession number: PRJNA268350). Bowtie2^92^ with default parameters (v2.5.4) was used for mapping fastq files to 771 PEP3458 family gene nucleotide sequences. Alignments with <95% identity or <100bp alignment length were discarded using msamtools (v1.1.3) and the number of reads mapping to each gene was counted using samtools (v1.23.1).

### Cell culture

J774 macrophages were cultured in DMEM (GIBCO) media supplemented with 10% heat-inactivated (HI-FBS) (Hyclone), penicillin (10,000 I.U./mL), streptomycin (10,000 ug/mL), and L-glutamine (2 mM). Any passaging and harvesting of J774 cells was performed by replacement of complete media and subsequent gentle scraping of cells using a double end, flat blade and J-hook Cell Lifter (Celltreat). J774 cells were cultured in 15-cm plates for screens, and either 10- cm plates or 6-well plates for flow cytometry assays or routine passaging of cells. HEK cells were maintained in DMEM, penicillin (10,000 I.U./mL), streptomycin (10,000 ug/mL), L-glutamine (2 mM), and FBS. Ramos cells were maintained in suspension culture in RPMI supplemented penicillin (10,000 I.U./mL), streptomycin (10,000 ug/mL), L-glutamine (2 mM), and 10% HI-FBS. All cells were maintained in a controlled humidified incubator at 37℃ and 5% CO2.

*Flow cytometry to measure polarization and cell surface expression in J774 macrophages* Flow cytometry on J774 macrophages exposure to purified cytokines were performed after 24 hours of stimulation with either IL-4 (50 ng/mL, PeproTech) or IFN**γ** (50ng/mL PeproTech) The following antibodies were used for J774 flow cytometry analysis: anti-mouse CD206 (MMR) (Biolegend, clone C068C2), anti-mouse CD80 (Biolegend, clone 16-10A1), anti-mouse Ly6a (Sca-1) (Biolegend, clone D7), anti-mouse IL-4 (Biolegend, clone 11B11), anti-FLAG antibody

(Biolegend, clone L5), anti-IgG Isotype Ctrl Armenian Hamster Monoclonal Antibody (Biolegend, clone HTK888). Antibodies were used at 0.5 µg per million cells. For flow cytometry assays with J774 macrophages, cells were harvested and centrifuged for 5 minutes at 500g, and resuspended in cell staining buffer (CSB) (BioLegend) containing TruStain Fcx PLUS (anti-mouse CD16/32) Antibody at a final concentration of 0.25 µg per 1×10^6^ cells (Biolegend) and incubated on ice for 15 minutes. Cells were subsequently centrifuged for 5 minutes at 500g at room temperature, resuspended in CSB containing the appropriate concentration of antibody, and incubated at 4℃ for one hour. Cells were subsequently washed once with CSB then resuspended in CSB for flow analysis. Negative antibody controls were run across all assays; additional IgG isotype controls were included for flow cytometry assays from co-culture experiments. Flow cytometry analysis was performed using Attune NxT Flow Cytometer (ThermoFisher) and analyzed using FlowJo software.

### Design of bacterial micropeptide library

Each bacterial micropeptide was codon optimized for mammalian expression using DNAchisel^93^ and two codon variants were generated for each given micropeptide, with the exception of a number of small microproteins where codon optimization constraints led to generation of only a single coding variant. An additional 250 random control micropeptide sequences were generated, leading to a final library size of 7,188 elements. In order to clone the library into the peptide display vector, primer binding sites and typeIIS BsmBI recognition sites were added to the 5′ and 3′ ends of each element. Each element in the library was normalized to an equal length of 260 nucleotides by inserting variable length DNA padding sequences between the primer binding and enzyme recognition sites.

### Cloning of bacterial micropeptide libraryh

The bacterial micropeptide library was synthesized by Twist Biosciences. First the library was amplified via PCR, with the following reactions established in PCR-free hood: 5 ng of template, 0.1 mL of each 100 mM primer, 1 mL of Herculase II polymerase, 1 mL of DMSO, 1 mL of 10 nM dNTPs, and 10 mL of 5x Herculase buffer. The following thermocycler conditions were used: 98℃ for 3 min, followed by 29 cycles at 98℃ for 20 s, 61℃ for 20 s, 72℃ for 30 s, and then a final extension step at 72℃ for 3 min. Amplified libraries were then run on a 2% TBE gel and extracted using manufacturer’s instructions (Qiagen). Libraries were then cloned into the lentivirus peptide display vector (MCB1174, addgene #228456), using a total of 4 × 10 mL GoldenGate reactions, which included 75 ng of pre-digested and gel-extracted backbone plasmid, 5 ng of amplified library (in a 2:1 molar ratio of insert to backbone), 0.13 mL of T4 DNA ligase (NEB, 20000 U/ml), 0.75 mL of Esp3I-HF (NEB), and 1 mL of 10x T4 DNA ligase buffer. These reactions underwent 30 cycles of digestion at 37℃ and ligation at 16℃ for 5 min each, followed by a final 5-min digestion at 37℃ and heat inactivation at 70℃ for 20 min. The resulting reactions were pooled and purified using MinElute columns (QIAgen), with an elution volume of 6 μL of ddH2O. Subsequently, 2 mL per tube was transformed into two tubes containing 50 mL of Endura electrocompetent cells (Lucigen) following the manufacturer’s instructions. After the recovery process, the cells were plated on 3-7 10′′ x 10′′ LB plates containing carbenicillin. Following overnight growth at 37℃, the bacterial colonies were scraped into a collection bottle, and plasmid pools were extracted using a HiSpeed Plasmid Maxiprep kit (QIAgen). To assess the quality of the libraries, 20 - 30 colonies were amplified from the plasmid pool for premium PCR (Primordium) to determine cloning efficiency and the proportion of empty backbone plasmids within the pools.

### Installation of peptide library into J774 macrophages

Large scale lentivirus production and infection of J774 macrophages was performed as follows: two T225 flasks were seeded with approximately 4×10^7^ HEK293 cells in 45mL of complete Optimem media and grown overnight to achieve roughly 75-85% confluency. The next morning, lipofectamine was used to transfect cells. For each flask, the following mixture was set up in a 15ml conical: 5mL of Optimem, 13μg pMD2.G, 30μg psPAX2, 42μg peptide library, 145μL lipofectamine P3000. In a second 15mL conical, 5mL of Optimem and 165μL of lipofectamine L3000 was added. The two mixtures were then combined and incubated for 15 minutes at room temperature. 20mL of media was removed from each flask, and then approximately 10ml of lipofectamine mixture was carefully added to cells, for a total of 35mL of volume per flask. Cells were incubated for six hours and transfection media was then removed and replaced with 45ml of warm cOptimem. 18 hours after media change, 45mL of viral supernatant was harvested. 45mL of warm cOptimem was added to cells, for a second viral supernatant harvest 24 hours after. The combined harvests were then centrifuged for 5 minutes at 500g and then viral supernatant was then filtered through a 0.45 μM PVDF filter (Millipore). Harvests were then concentrated at a ratio of 1:3 of Lenti-X (Takara Bio) to viral supernatant and incubated at 4C for approximately 6 hours. The mixture was centrifuged for 45 minutes at 1,500g. Supernatant was discarded and pellet was resuspended at 1000x concentration in DMEM. All of the concentrated supernatant was resuspended into complete DMEM and used to infect approximately 1×10^8^ J774 macrophages across eight 15cm plates, in order to infect at >1000x coverage the number of library elements. Lentivirus containing media was maintained on cells for 48 hours, and afterwards flow cytometry was used to measure percentage of mCherry positive cells as a proxy for transduction efficiency. Lentivirus was then removed and the library was expanded for subsequent cell sorting.

### Library flow cytometry and cell sorting

For the sorting based screen, J774 macrophages were maintained at >1000x library coverage throughout. For each replicate, approximately 1×10^8^ cells were harvested (15-20% of those cells estimated to have a peptide library based on percentage of mCherry positive cells via flow cytometry. All centrifuge steps were at 500g for 5 minutes at room temperature and volume of cell staining buffer (CSB) (BioLegend) was maintained at 1ml per 1×10^7^ cells throughout.

Harvested cells were centrifuged, media was aspirated and cells were resuspended in 1ml per 1×10^7^ cells of CSB. Cell pellets were then centrifuged and resuspended in CSB containing TruStain Fcx PLUS (anti-mouse CD16/32) Antibody at a final concentration of 0.25 µg per 1×10^6^ cells and incubated for 15 minutes on ice. Pellets were then centrifuged and resuspended in CSB containing 0.5 µg per million cells of anti-mouse CD80 or anti-mouse Ly6a antibody and placed in the dark at 4℃ for one hour. Pellets were washed twice with CSB and resuspended at 1ml per 1×10^7^ CSB containing 1mM EDTA and filtered through 5ml round-bottom tubes with cell strainer (Corning). Sorting was performed on an instrument in the Shared FACS Facility obtained using NIH S10 Shared Instrument Grant (FACSAriaII: S10RR025518-01, FACSAria fusion: Purchased by Parker Institute for Cancer Immunotherapy). Cells were sorting into 15ml conicals containing 10% FBS. After completion of sorting, cells were centrifuged, supernatant was aspirated and cells were stored at -80C for subsequent genomic DNA extraction.

### Library preparation and sequencing

For genomic extractions of library samples, the QiaAmp Mini Kit was used (Qiagen). First, cell pellets were resuspended in 200μL of PBS. Then 20μL of protease (Qiagen) and 200μL of AL buffer was added to each sample, which were vortexed for 10-15 seconds and then incubated at 56℃ for 10 minutes. Then 200μL of ethanol was added, samples were vortexed for 10-15 seconds and centrifuged through the provided column for 30 seconds at 8000 rpm. 500μL of AW1 buffer was added, samples were centrifuged for the 30 seconds at 8000 rpm. 500μL of AW2 buffer was added and samples were centrifuged for 30 seconds at 13000 rpm. The columns were then centrifuged without buffer for another three minutes at 13000 rpm. Columns were placed into a fresh microcentrifuge tube and 50μL of EB was added to columns and incubated at room temperature for five minutes. Samples were then centrifuged for one minute at 8000rpm and quantified using a nanodrop. Library amplification was performed using a modified version of the HT-recruit protocol^44^. 6-24x 25μL PCR reactions were set up on ice (in a clean PCR hood to avoid amplifying contaminating DNA), with the number of reactions depending on the amount of genomic DNA available in each experiment. 500ng of genomic DNA, 0.125 mL of each 100 mM primer, and 12.5μL of NEBnext 2x Master Mix (NEB) was used for each reaction. After optimization of cycle number and genomic DNA quantity, the following thermocycler conditions were used: 3 minutes at 98℃, then 28x cycles of: 98℃ for 10 s, 63℃ for 30s, 72℃ for 30s, then a final step of 72℃ for 2 minutes. PCR reactions were then pooled and at least 150μL of PCR product was run on a 2% TBE gel for at least one hour. Gel bands were extracted and purified using QIAquick Gel Extraction kit (QIAgen), and eluted in 30 μL of EB. Libraries were then quantified using the Qubit HS kit (ThermoFisher). Paired-end sequencing (150 bp) was performed by Novogene on the Novaseq X Plus platform, with a minimum of 7.5Gb of data per sample.

### Bioinformatic analysis of microprotein screen

For sequencing analysis of peptide display screen, removal of adapter sequences was performed using AdapterRemoval (v2.3.4)^94^. Subsequently, constant regions surrounding peptides of interest were removed using trimmomatic (v0.39)^95^. Mapping of peptides was performed using prior HT-recruit analysis^44^. In brief, a reference containing all peptides in the library was generated using the script ‘makeIndices.py’ and reads were aligned with 0 mismatch allowance using the script ‘makeCounts.py’. The relative enrichment of each peptide in the top 10% (high) of either ly6a positive or CD80 positive relative to the bottom 10% (low) of each population was computed using the ‘makeRhos.py’ script. To remove low-confidence measurements, microproteins with low counts were removed from the analysis. We chose to exclude library elements with five or fewer counts in either sorting bin (high or low) from either experimental replicate. For each library element, scores were calculated as log2((HighCount/TotalHigh):(LowCount/TotalLow)). For hit calling, microproteins were required to have both variants and both replicates exceed two standard deviations above the mean log2 score of the random negative controls.

### Generation of single peptide vector constructs

For expression of single micropeptides in the display system, either Geneblocks or oligonucleotides were synthesized (IDT) containing BsmBI cut sites. Oligonucleotides were annealed by incubating 5 μM of each oligo in 2x annealing buffer (200mM Potassium Acetate, 60 mM HEPES-KOH pH 7.4, 4 mM Magnesium Acetate) under the following thermocycler conditions: 95℃ for 5 min, cool to 22℃ at ramp rate 0.1℃/s, 22℃ for 2 min. 100 ng of the respective vector was incubated at a 3:1 molar ratio with 10x T4 ligase buffer and Golden Gate Assembly Kit (BsmBI-v2) (NEB), at following thermocycler conditions: 42℃ for 60 minutes, 60℃ for 5 minutes. 2μl of each product was then mixed with 50μl of DH5 competent *E. coli* heat-shocked at 42℃ for 30 seconds, then incubated at 37℃ for approximately 30 minutes supplemented with SOS buffer (Fisher), then selected on LB carbenicillin plates. Single colonies were grown in LB containing a working concentration of 50μg/ml of carbenicillin, and harvested for plasmid purification per manufacturer’s instructions (Qiagen). All plasmids were confirmed via whole plasmid sequencing (Plasmidsaurus).

### Transduction of individual display vectors into J774 macrophages

First lentivirus was produced by incubating 1000ng of display vector, 750ng of psPAX2 and 250ng pMD2G with 13μL of polyethylenimine (PEI) was added to 200μl of serum free DMEM and incubated for approximately 15 minutes. The mixture was then added to approximately 1×10^6^ HEK293 cells. Viral supernatants were harvested at 48 hours and 72 hours, and media was combined and filtered through a 0.45 μM SFCA filter. The filtered supernatant was then used to infect approximately 0.5×10^6^ J774 macrophages. Lentivirus containing media was maintained on cells a minimum of 48 hours with and afterwards replaced with complete DMEM for 24 hours. For indicated assays, LPS (1 ng/mL) was added 24 hours prior to flow cytometry experiments and/or RNA extraction.

### RNA-seq and qRT-PCR experiments

For transcriptional profiling of J774 macrophages, 1×10^6^ cells were plated in 6-well plates. For samples stimulated with either IFN**γ** or IL-4, 50 ng/mL was added to the media 24 hours prior to harvesting RNA. Media was then aspirated and for extraction of RNA, 500ul of trizol was added to each well, and incubated at room temperature for 5 minutes. Samples were then transferred to Eppendorf tubes and 100ul of chloroform was added. Samples were vigorously vortexed and incubated at room temperature for 2 minutes, then centrifuged at 4℃ at 15000rpm for 15 minutes. The upper phase of the supernatant was transferred to a new Eppendorf tube, and 200ul of 70% ethanol was added. Samples were then run through the RNeasy Mini spin column (Qiagen), per manufacturer’s instructions. On-column DNA digestion was performed, using 10μL Qiagen DNase I with 70μL Buffer RDD per sample and 30 minutes of incubation at room temperature. Samples were eluted in 50μL RNase free water and purified RNA was sent to Novogene for mRNA library preparation (polyA enrichment) and sequencing (Novaseq 6000, 150bp paired end reads, minimum of 6Gb raw reads per sample). For qRT-PCR experiments with J774 macrophages, RT was performed using superscript and cDNA amplified using SYBR green master mix (BioRad), both via manufacturer’s instructions. TaqMan PCR was used for gene expression of iPSC macrophages, per manufacturer’s instructions. All qPCR was performed using the CFX384 Real-Time PCR System (BioRad). The ΔΔCt method was used for all expression analysis.

### RNA-seq analyses and differential expression analysis

Fastq files were mapped using STAR (v2.7.11b)^96^ and gene-level counts were generated using HTSeq (v2.1.2)^97^. Differential gene expression analysis was performed using DESeq2 (v1.50.2) in R^98^. Wald test statistics were used to identify differentially expressed genes (DEGs). Genes with a Benjamini-Hochberg adjusted p-value < 0.05 and |log2 fold change| > 1 were considered statistically significant unless otherwise noted. Pathway enrichment analysis was performed using clusterProfiler with Gene Ontology Biological Process (GO BP) and KEGG databases for over-representation analysis (ORA), and the fgsea package (v1.36.2) for gene set enrichment analysis (GSEA) against the MSigDB Hallmark gene set collection^99^. Semantic similarity reduction of GO terms was performed using the simplify() function with a cutoff of 0.4.

### Phagocytosis assays

J774 macrophages were lentivirally transduced with indicated peptide display constructs, and subsequently seeded into a 24-well plate at a density of 50k/well. After 24hours, LPS was added at a final concentration of 100ng/mL for another 24 hours. To establish the co-culture assays, WT Ramos cells were harvested and centrifuged at 500g for 3 minutes, washed once with 1x PBS, and resuspended in PBS with pHrodo-Red succinimidyl ester (ThermoScientific), at a final dilution of 1:50000 from the stock concentration, and Calcein AM (for cancer live cell staining) (ThermoFisher) at final dilution of 1:3000 from the stock concentration. Ramos cells were then incubated for 15 minutes, then centrifuged at 500g for 3 minutes. Cells were then washed with DMEM containing HI-FBS once, and then resuspended with DMEM (+HI-FBS) containing anti-CD20 at 15ug/mL (Rituxan) and anti-CD47 antibody at 10ug/mL (clone B6.H12, BioXCell). The pHrodo red labeled Ramos cells were incubated in this media for 10 minutes, then 250k cells per well were added to each well containing J774 macrophages. Plates were transferred to an incubator and imaged every 60 min using an Incucyte (Essen). Total red intensity for each well, averaged over 9–16 images per well, was calculated after applying a threshold for red intensity that excluded most or all cells at the first time point using top-hat background subtraction. Reported values represent the mean total red fluorescence intensity at each timepoint.

### Bacterial strains and growth conditions

All bacterial strains and plasmids are listed in Supplementary Table 10. ClearColi (Fisher) cells were inoculated from glycerol stocks and grown overnight at 37 °C in 5 ml Luria broth (LB). The next day, the OD_600 nm_ of cultures was measured and diluted to OD_600 nm_ = 0.1 in 5 ml of fresh LB. Cells were harvested for downstream assays in the exponential growth phase after a 2.5-3 h incubation at 37 °C. Antibiotics for plasmid maintenance were used at the following concentrations: spectinomycin (100 ug/mL), carbenicillin (100 ug/mL). *Leptotrichia* sp. oral taxon 879 str. F0557 was routinely cultured on Brain Heart Infusion Blood (BHIB) agar (Hardy Diagnostics) at 37°C anaerobically (90% nitrogen, 5% carbon dioxide, 5% hydrogen) in an anaerobic chamber (Sheldon Manufacturing) for 48 h.

### Leptotrichia whole genome sequencing

*Leptotrichia* str. F0557 was grown as described above. Cell biomass was harvested by scraping a BHIB agar plate, followed by resuspension and washing in PBS, and final resuspension in DNA/RNA Shield (Zymo). Genomic sequencing was performed with Plasmidsaurus.

### Western blot analysis

To probe for micropeptide expression, ClearColi strains were grown overnight to saturation, subcultured to OD_600_ = 0.1, and incubated for 3h. Cell density was measured with turbidimetry and cell counts were normalized, then aliquots of each strain were pelleted at 15,000xg for 3 min at 4°C. The supernatant was decanted and cell pellets were flash frozen in liquid nitrogen. Pellets were subsequently thawed and resuspended in NuPAGE LDS Sample Buffer (Fisher) supplemented with 50 mM dithiothreitol (DTT). Each sample was boiled at 95°C for 10 min and run on a 12% NuPAGE Bis-Tris gel (Fisher) prior to transfer to a 0.2-μm nitrocellulose membrane. HRP Anti-6X His antibody (Abcam, catalogue no. ab1187) was used at a final concentration of 1:5000, and SuperSignal West Pico PLUS Substrate (ThermoFisher) was used to develop the blots. Blots were imaged using an iBright 750 Imaging system (Invitrogen).

### Co-culture assays with ClearColi

J774 macrophages were seeded into a 6-well plate at a density of 250k/well. ClearColi strains were grown overnight to saturation, subcultured to OD_600_ = 0.1, and incubated for 3h. ClearColi cells were pelleted at 15,000xg for 3 min, washed once with PBS, and cell density was measured with turbidimetry. ClearColi cells were added to the J7 cells at an MOI of 10 in DMEM (+HI-FBS) and plates were then centrifuged for 3 minutes at 300g prior to incubation. ClearColi colony forming units (CFUs) were quantified for each co-culture assay at the start and end of each experiment. At the indicated time points, co-cultured cells were harvested for either flow cytometry or RNA extraction after overnight incubation. For RNA-seq, RNA extraction was performed using the RNAeasy Mini Spin Column (Qiagen), per manufacturer’s instructions. On-column DNA digestion was performed, using 10μL Qiagen DNase I with 70μL Buffer RDD per sample and 30 minutes of incubation at room temperature. Samples were eluted in 30μL RNase free water. Purified RNA was sent to Novogene for mRNA library preparation (polyA enrichment) and sequencing (Novaseq 6000, 150bp paired end reads, minimum of 6Gb raw reads per sample).

### Data visualization

Initial plots were generated by Python (Plotly) and R (ggplot) code. Plots were organized and refined with Adobe Illustrator (v29.6.1).

## Data Availability

Microprotein library screen short read sequencing is available under NCBI BioProject ID PRJNA ***. Whole genome sequencing raw data for *Leptotrichia* str. F0557 are available NCBI BioProject ID ***.

## Figures

**Extended Data 1.**
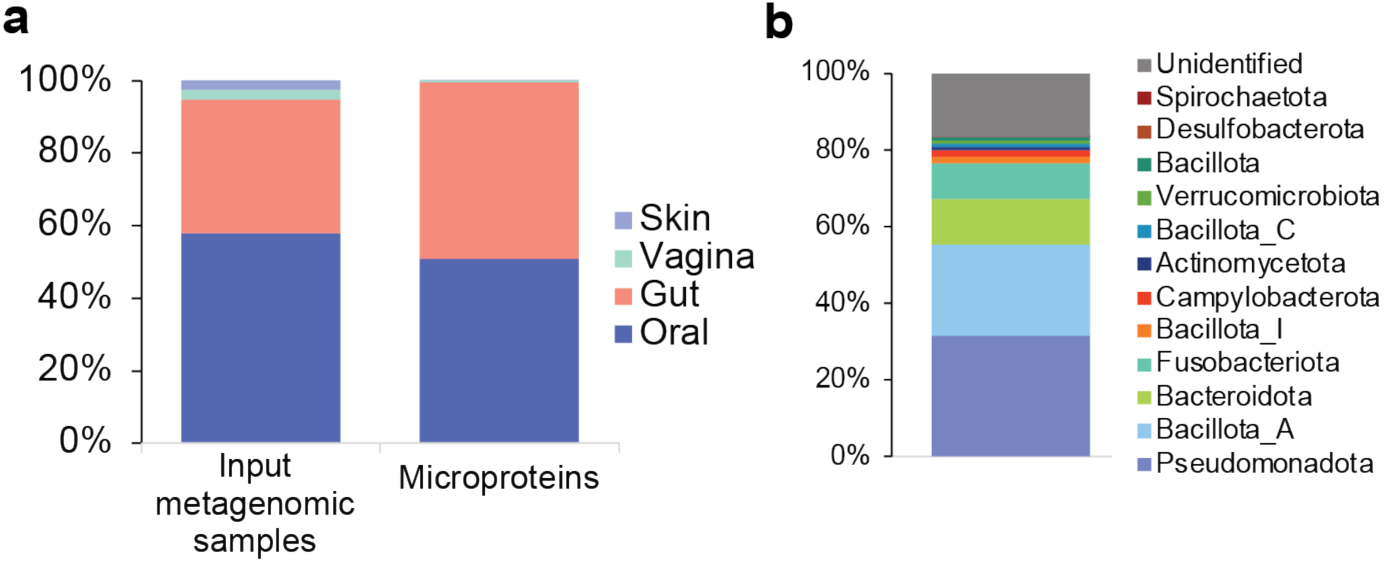
Characteristics of synthesized microprotein library. a. Comparison between the anatomic distribution of the input metagenomic samples used and the distribution of microbial microproteins included in the final library b. Distribution at the phylum-level of microbial microproteins included in the final library

**Extended Data 2.**
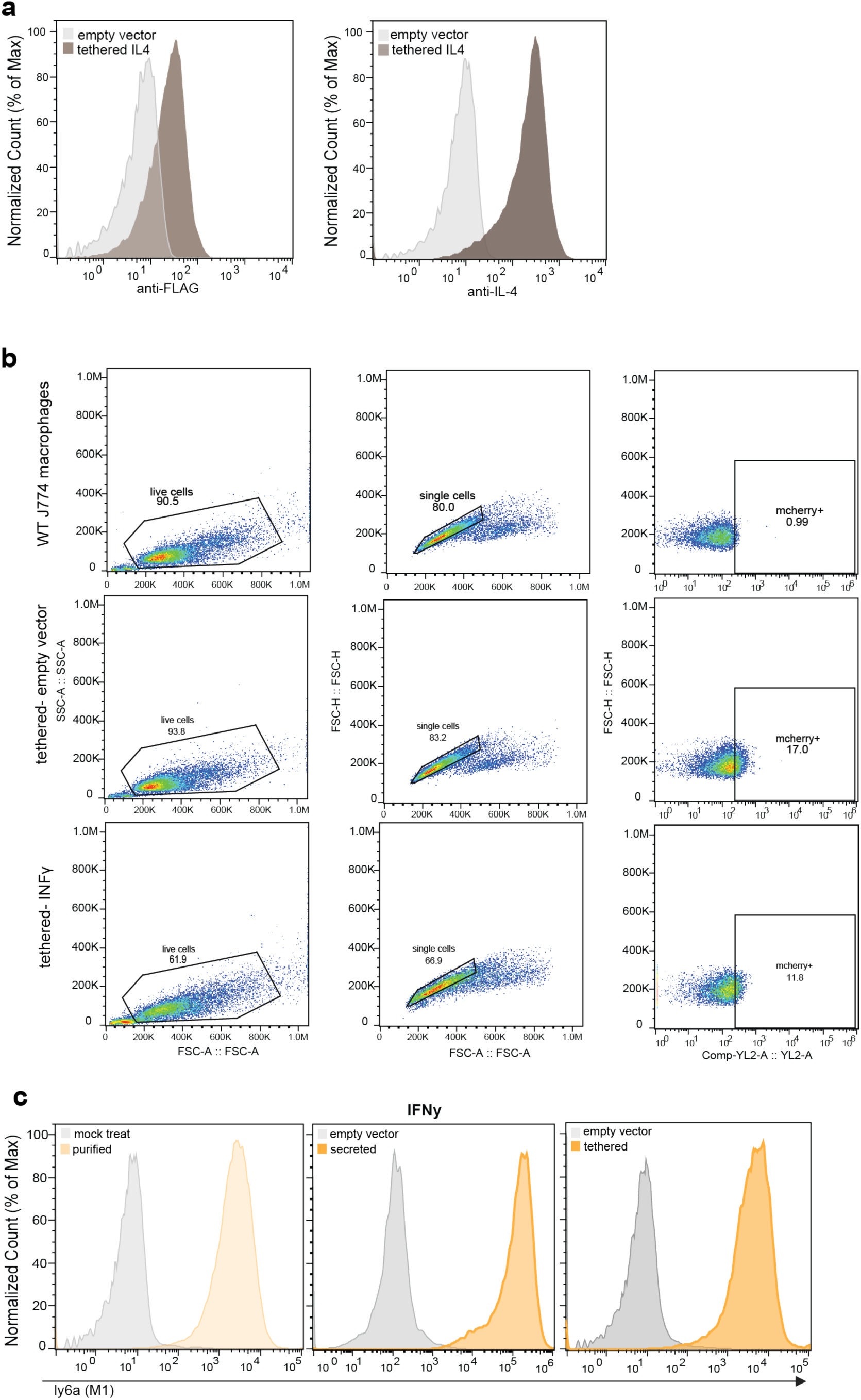
Display of cytokines in macrophages is sufficient to polarize macrophages to distinct cell states. a. Confirmation of cell surface expression utilizing peptide display system. J774 macrophages expressing tethered IL-4 along with empty vector control were stained with either anti-FLAG or anti-IL4 antibody. b. Representative gating strategies shown for peptide display constructs. c. Flow cytometry histogram plots comparing Ly6a expression of J774 macrophages expressing tethered or secreted IFN**γ**, as well as exposed to purified IFN**γ** (orange), relative to negative controls (grey).

**Extended Data 3.**
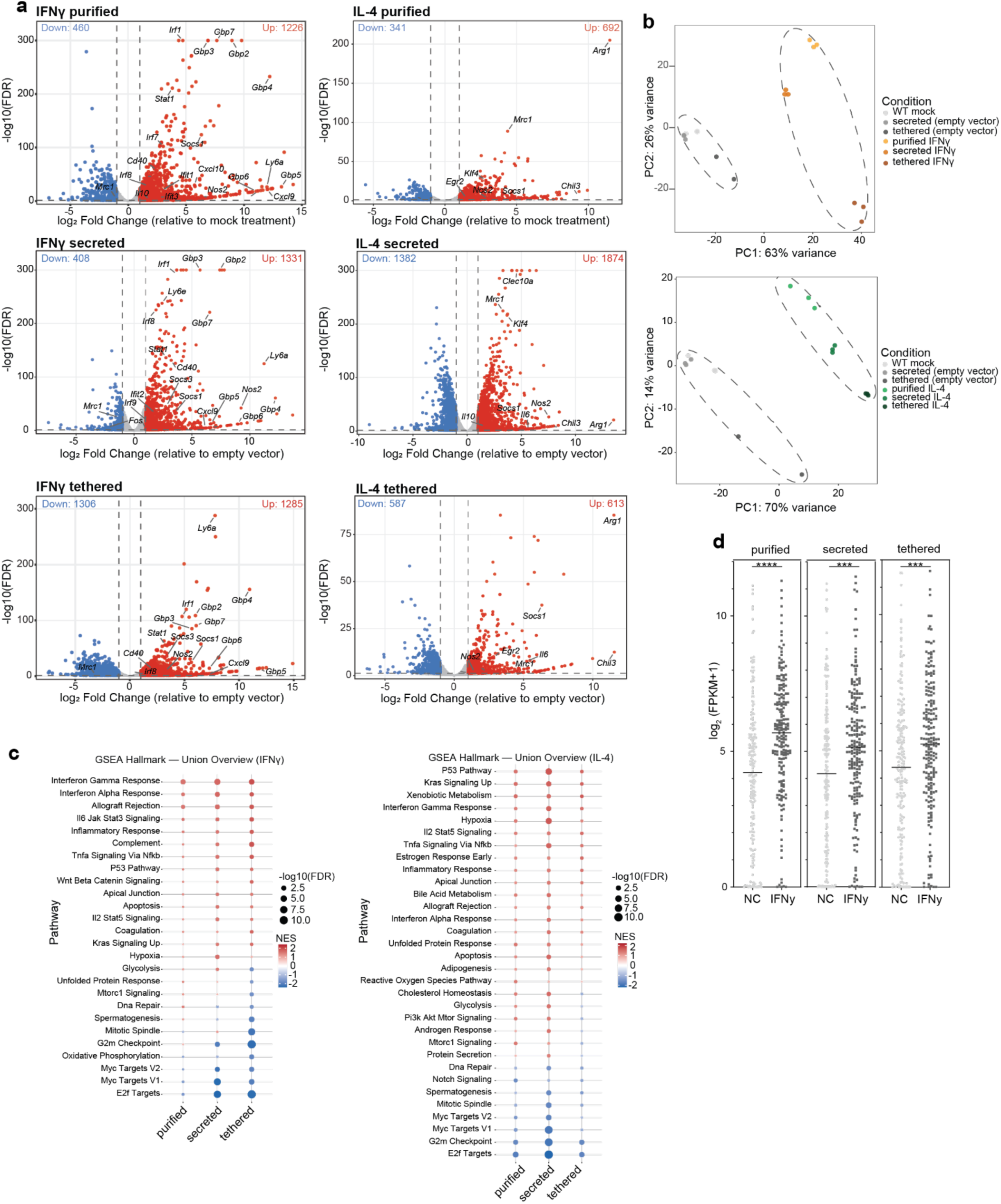
Shared transcriptional profile between exogenous cytokine exposure and cytokine display expression in J774 macrophages. a. Bulk RNA-seq of J774 macrophages expressing either tethered or secreted cytokines, as well as purified cytokine controls. n=2 distinct biological samples for tethered empty vector control, and n=3 for all other samples. Red dots represent genes differentially upregulated, blue dots represent genes differentially downregulated, relative to respective negative control. Select individual genes associated with M1 or M2 polarization are labeled. b. Principal component analysis plots of bulk RNA-seq demonstrating clustering of macrophages expressing either IFN**γ** via the surface display systems (green) or IL-4 (yellow) and each purified cytokines, relative to negative controls. c. Gene set enrichment analysis of IFN**γ** (left) and IL-4 (right) induced transcriptional programs across Purified, Secreted, and Tethered stimulation conditions. Each dot represents a GO Biological Process pathway. Dot color indicates normalized enrichment score (NES), with red indicating positive enrichment and blue indicating negative enrichment. Dot size corresponds to –log10(FDR). d. Analysis of RNA-seq data comparing transcriptional changes of individual genes previously associated with IFN**γ** stimulation. Gene list was generated from individually upregulated genes via scRNAseq from BMDM macrophages exposed to IFN**γ** stimulation (Cui et al., 2024). Unpaired t-test performed to determine statistical significance.

**Extended Data 4.**
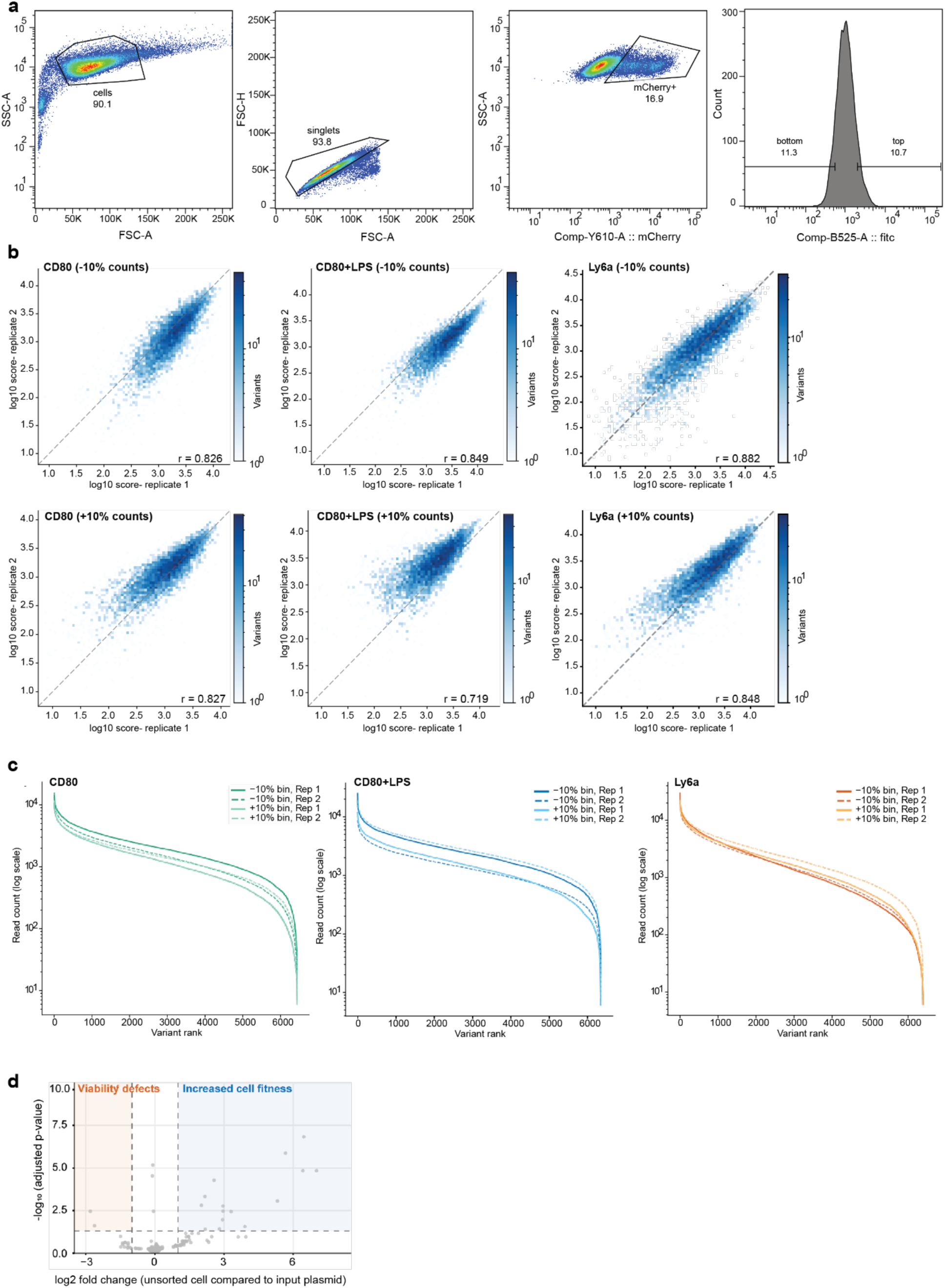
Screening approach to identify immunomodulatory peptides. a. Representative gating strategy of peptide display screen. J774 macrophages were first gated for the presence of mCherry. mCherry positive cells were then sorted into either high (CD80 or Ly6a) or low (CD80 or Ly6a) antibody signal. Appropriate compensation controls were performed for all samples. b. Replicate concordance of sorted fractions counts across peptide display screens. Log10 read counts are plotted for Replicate 1 versus Replicate 2 for the −10% (top) +10% (bottom) sorted fractions across all three screens (left to right): CD80, CD80+LPS, Ly6a. Pearson correlation coefficients and variant counts are shown for each panel. c. Count distributions for all conditions of each individual sample from each respective peptide display screen performed. d. Volcano plot of DESeq2 results comparing the unsorted bulk population of transduced J774 macrophages to the plasmid input library. Each point represents one coding variant. The x-axis shows the apeglm-shrunken log2 fold-change; the y-axis shows −log10 adjusted p-value. Vertical and horizontal dashed lines indicate thresholds of log2FC = −1 and padj = 0.05, respectively. The orange shaded region marks the intersection of both thresholds for candidate microproteins with viability defects, and the blue region marks the threshold for candidate microproteins with pro-growth phenotypes

**Extended Data 5.**
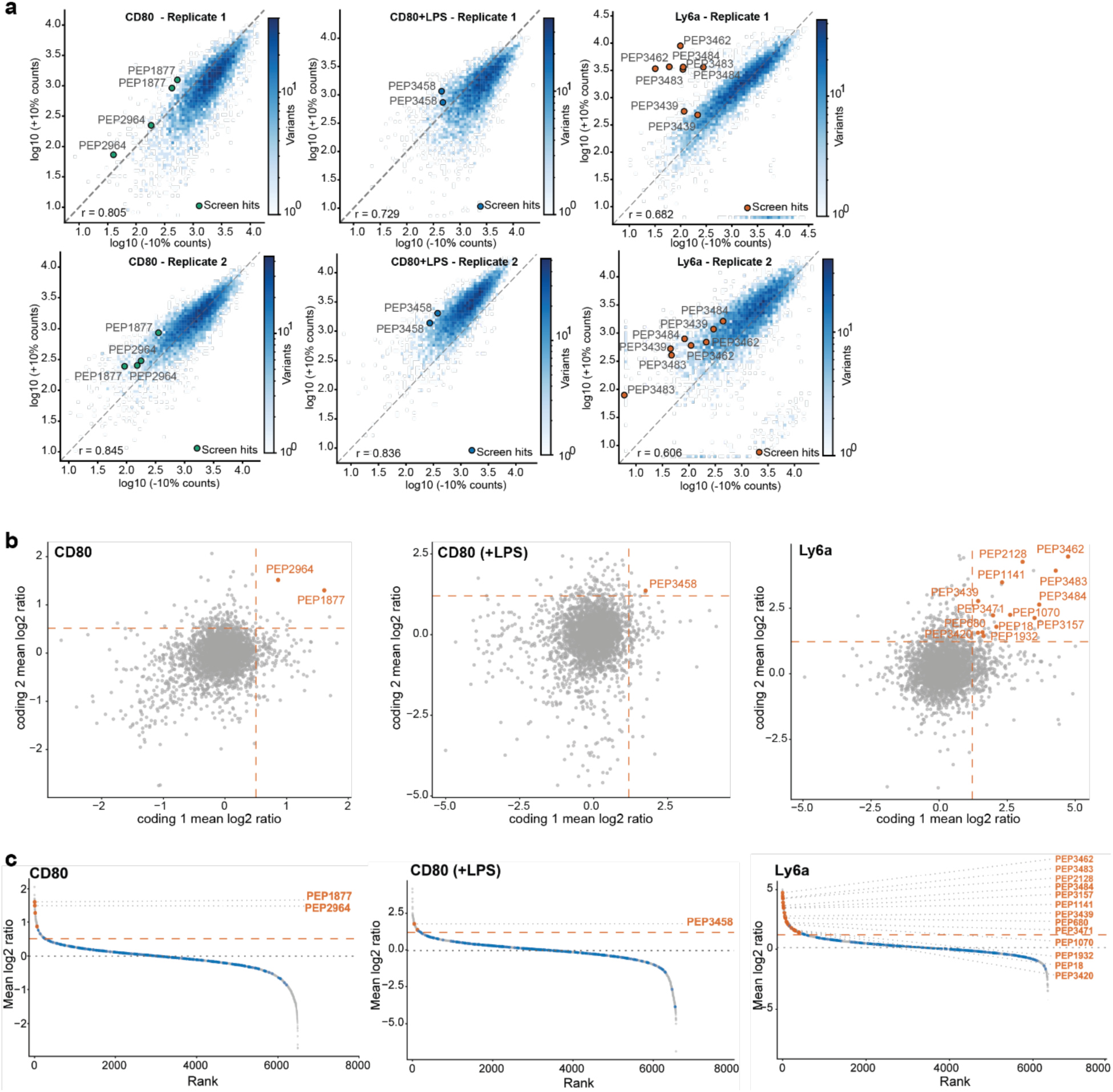
A high-throughput screen identifies candidate immunomodulatory microproteins. a. Log10 transformed read counts from the low (bottom 10%) and high (top 10%) sorted fractions plotted separately for each biological duplicate (top: replicate 1, bottom: replicate 2) across all the three screens (from left to right): CD80, CD80+LPS, Ly6a. Each point represents one individual coding variant of the microproteins present in the library. Pearson correlation coefficients and variant counts are shown for each panel. Screen hits are highlighted as colored points. b. Scatter plot showing average mean log2 ratio of biological replicates. Each point represents one peptide base plotted as the mean log2 ratio of coding variant 1 against coding variant 2. Candidate microprotein hits are labelled in orange. Dashed orange lines indicate the hit threshold on each axis, with threshold calling as previously described. c. Ranked plot microproteins, ranked in descending order of mean log_2_ ratio. The hit threshold (dashed orange line) was derived from the random control distribution as previously described. Hit labels are shown at the right margin.

**Extended Data 6.**
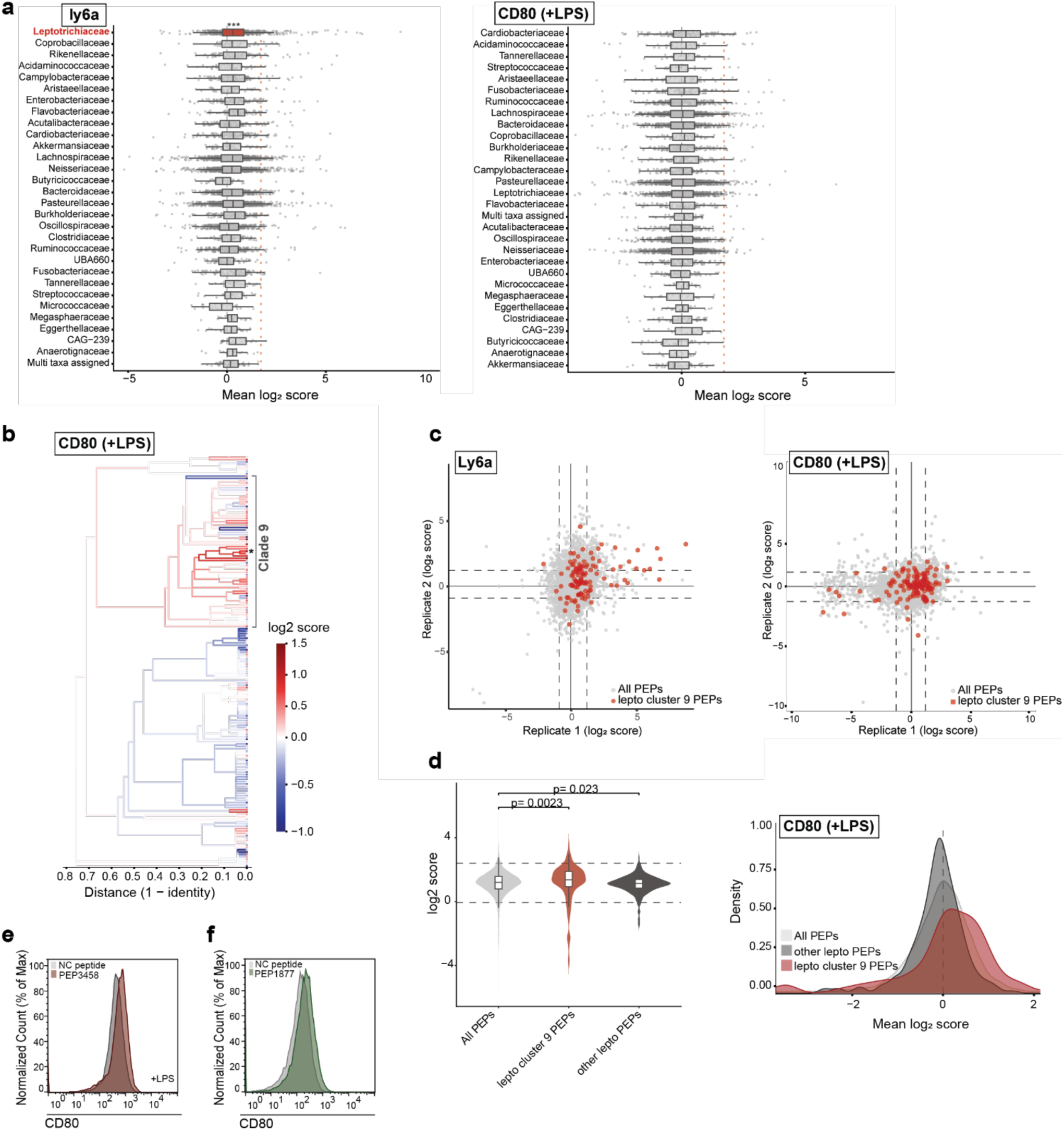
*Leptotrichia* Contain cluster of microproteins with putative immunomodulatory activity. a. Horizontal boxplot showing the distribution of mean log₂ scores for bacterial peptide families in the Ly6a (left) and CD80+LPS (right) macrophage polarization screen. Each row represents one bacterial family. The x-axis shows the screen score mean log₂ score. Families are ranked from top to bottom in descending order of the percentage of outlier peptides. Only families with at least 20 score observations after count filtering are shown. Grey points show individual score observations (biological replicates treated as independent observations). The dotted orange vertical line indicates the outlier threshold, defined as the background mean + 2 SD computed from the full pre-join screen distribution. Boxes and axis labels are colored red for families that are significantly enriched for outlier peptides (BH-adjusted p < 0.05, one-sided Fisher’s exact test); non-significant families are shown in grey. Peptides without a family assignment, including human-derived peptides, were excluded. b. Sequence-based hierarchical clustering of 175 Leptotrichia peptides colored by the CD80(+LPS) macrophage polarization screen scores. Hierarchical clustering was performed for all peptide sequences using single-linkage (nearest-neighbor) agglomerative clustering on a pairwise sequence identity distance matrix. Sequence identity between each pair of peptides was computed using a sliding-window best-match approach. Dendrogram branches are colored by the mean CD80(+LPS) log₂ score of all leaf nodes below that branch, and each leaf dot is colored by the mean CD80(+LPS) score for both biological replicates. Leaf dots with asterisks represent screen hits. c. Scatter plot with log₂ enrichment scores for all microproteins in the Ly6a and CD80(+LPS) screen, represented as an XY scatter plot reflecting biological replicates. Grey points represent all screened microproteins; *Leptotrichia* clade 9 family microproteins labeled in red. Dashed lines indicate ±2 standard deviations from the mean of random negative control peptides. d. Violin plot (left) and density plot (right) of mean log2 scores (average of biological replicates) per microprotein for each group indicated. *Leptotrichia* clade 9 microproteins indicated in red, *Leptotrichia* peptides from all other clades represented in dark grey. All other microproteins (excluding the *Lepotrichia* microproteins) are represented in light grey. Statistical comparisons were performed between each *Leptotrichia* group and the all peptides background using a Mann-Whitney U test. Exact p-values are shown on each comparison bracket. e. Flow cytometry histogram plots demonstrating increased CD80 expression in J774 macrophages expressing *Leptotrichia* PEP3458 (red), in comparison to negative control peptide (in grey) f. Flow cytometry histogram plots demonstrating increased CD80 expression in J774 macrophages expressing *Coprobacillus* PEP1877 (green), in comparison to empty vector and negative control peptides (in grey)

**Extended Data 7.**
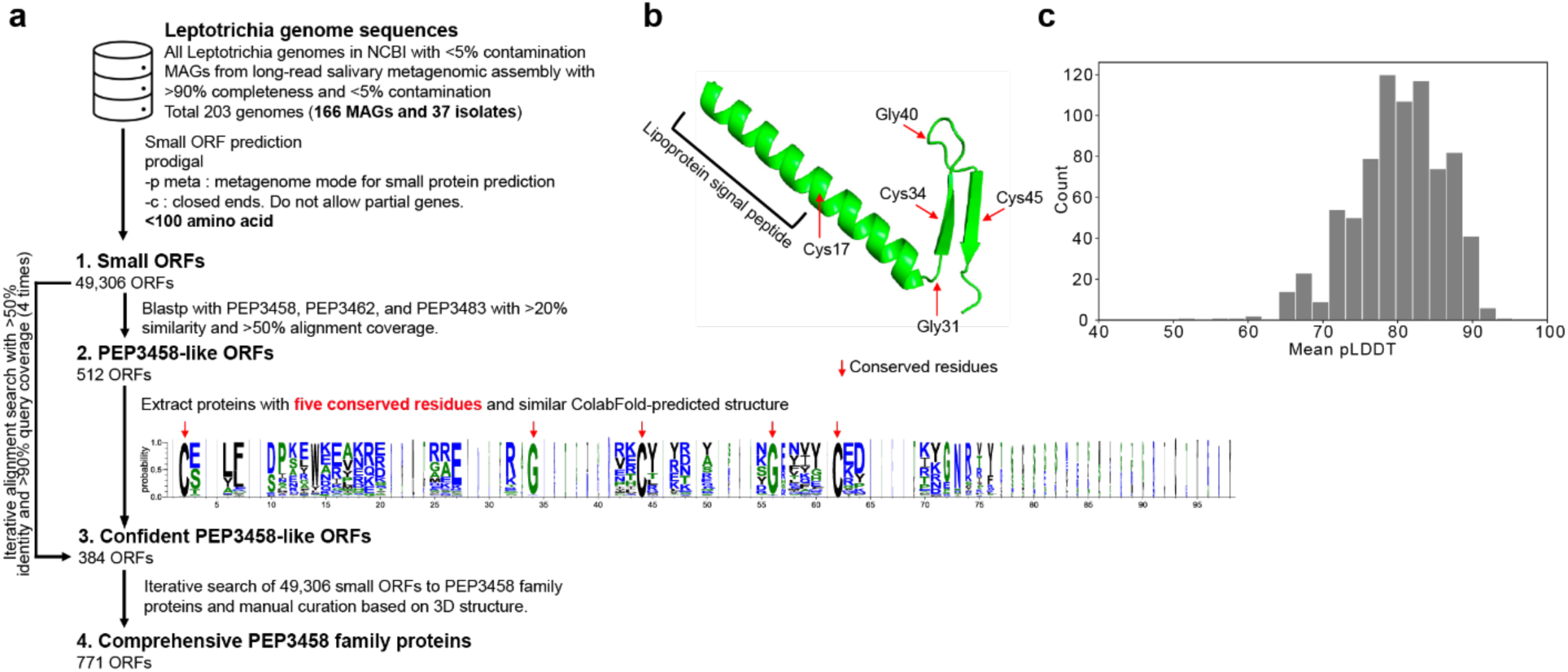
Generating a comprehensive collection of PEP3458 family microproteins from *Leptotrichia* genomes. a. Computational workflow for collecting PEP3458 family microproteins. The genome collection comprised 166 high-quality MAGs and 37 isolated genomes. Small ORFs (<100 aa) were predicted with Prodigal, yielding 49,306 ORFs, which were searched by BLASTP against the macrophage pro-inflammatory proteins PEP3458, PEP3462, and PEP3483 with >20% identity and >50% query coverage, giving 512 PEP3458-like ORFs. From these, 384 ORFs possessing the five conserved residues identified by multiple sequence alignment and a ColabFold-predicted structure similar to PEP3458 were defined as confident PEP3458-like ORFs. The sequence logo shows residue conservation across this set; red arrows indicate the five conserved residues. We then iteratively searched all 49,306 small ORFs against the confident set with >50% identity and >50% query coverage, adding new hits as references each round until convergence (4 iterations). Candidates with predicted structures clearly divergent from PEP3458 were excluded by manual curation, yielding 771 PEP3458 family proteins. b. ColabFold-predicted structure of PEP3458 protein. The lipoprotein signal peptide region located at the N-terminus of the protein. Five conserved residues in this family protein are shown as red arrows. c. Distribution of mean pLDDT (ColabFold confidence score) for the 771 PEP3458 family proteins.

**Extended Data 8.**
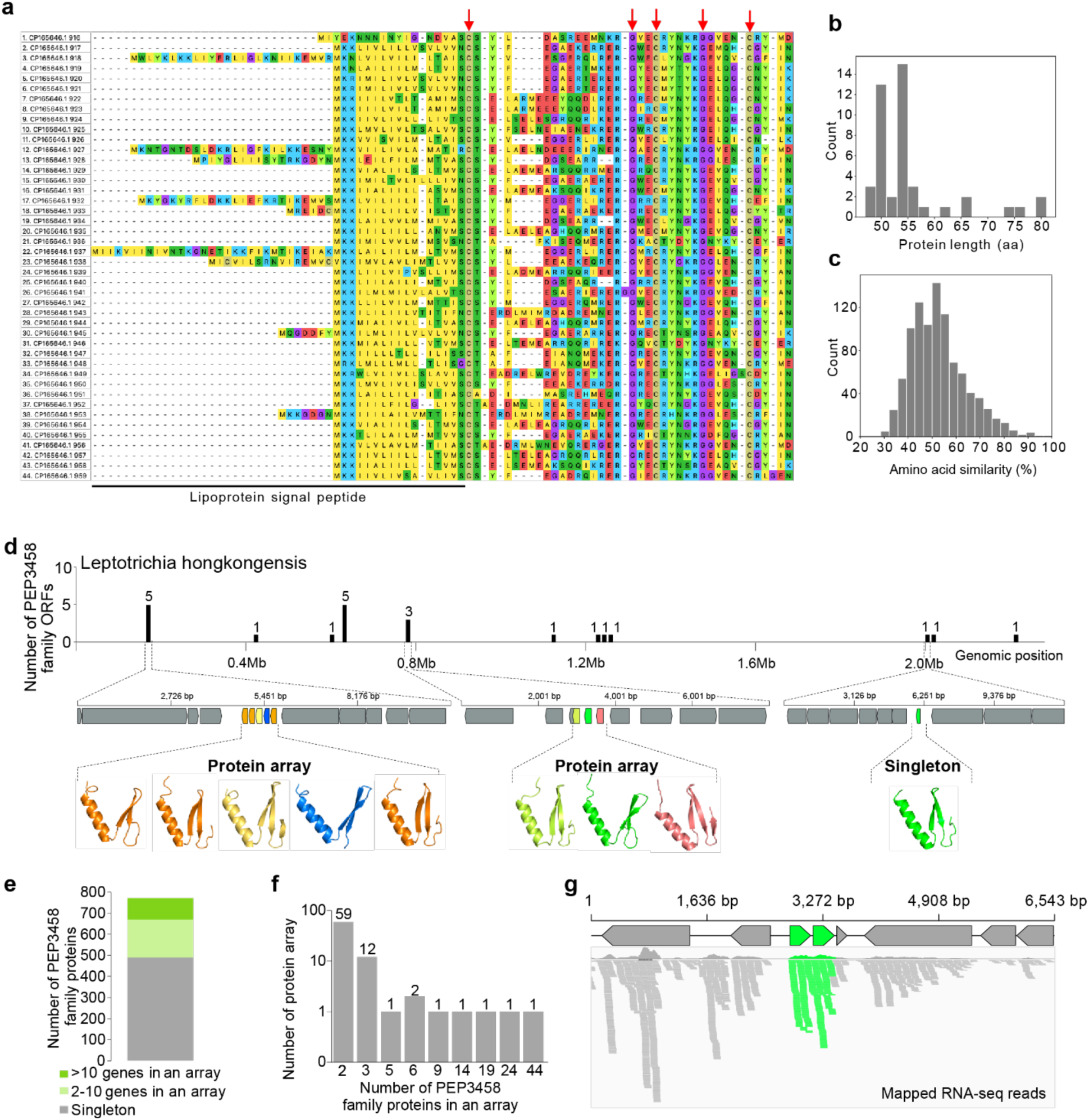
Genetic characterization of *Leptotrichia* protein array. a. Multiple sequence alignment of the 44 PEP3458 family ORFs in the protein array. The alignment was obtained using MUSCLE. The lipoprotein signal peptide region is indicated by a black line below the alignment; red arrows above mark the five conserved residues. b. Distribution of protein lengths within the array shown in (a). c. Distribution of pairwise amino acid identity (%) obtained using BLASTP within the array shown in (a) and main Fig. 3d. d. Representative genome (*L. hongkongensis*) carrying mini protein arrays. The x-axis indicates genomic position; the y-axis shows the number of PEP3458 family ORFs per 15 kb window. Genetic maps below show two representative mini-array loci and one singleton locus; coordinates in bp. PEP3458 family ORFs are colored by structural clade as in main Fig. 3a, and flanking genes are gray. e. The proportion of PEP3458 family ORFs encoded in the protein array. The bar shows the fraction of PEP3458 family ORFs found as singletons, in mini arrays (2–10 ORFs), and in large arrays (>10 ORFs), summed across all Leptotrichia genomes. f. The distribution of the number of PEP3458 family ORFs in a protein array. The x-axis shows the number of PEP3458 family ORFs in a protein array, and the y-axis indicates the number of protein arrays. g. Representative transcriptional activity of a PEP3458 family array in the human oral cavity. PEP3458 family ORFs are shown as green arrows and flanking genes as gray arrows. The amino acid similarity is 72% between the two PEP3458 family proteins. Bars below show mapped read coverage; green bars indicate reads mapped to PEP3458 ORFs and gray bars indicate reads mapped to flanking genes.

**Extended Data 9.**
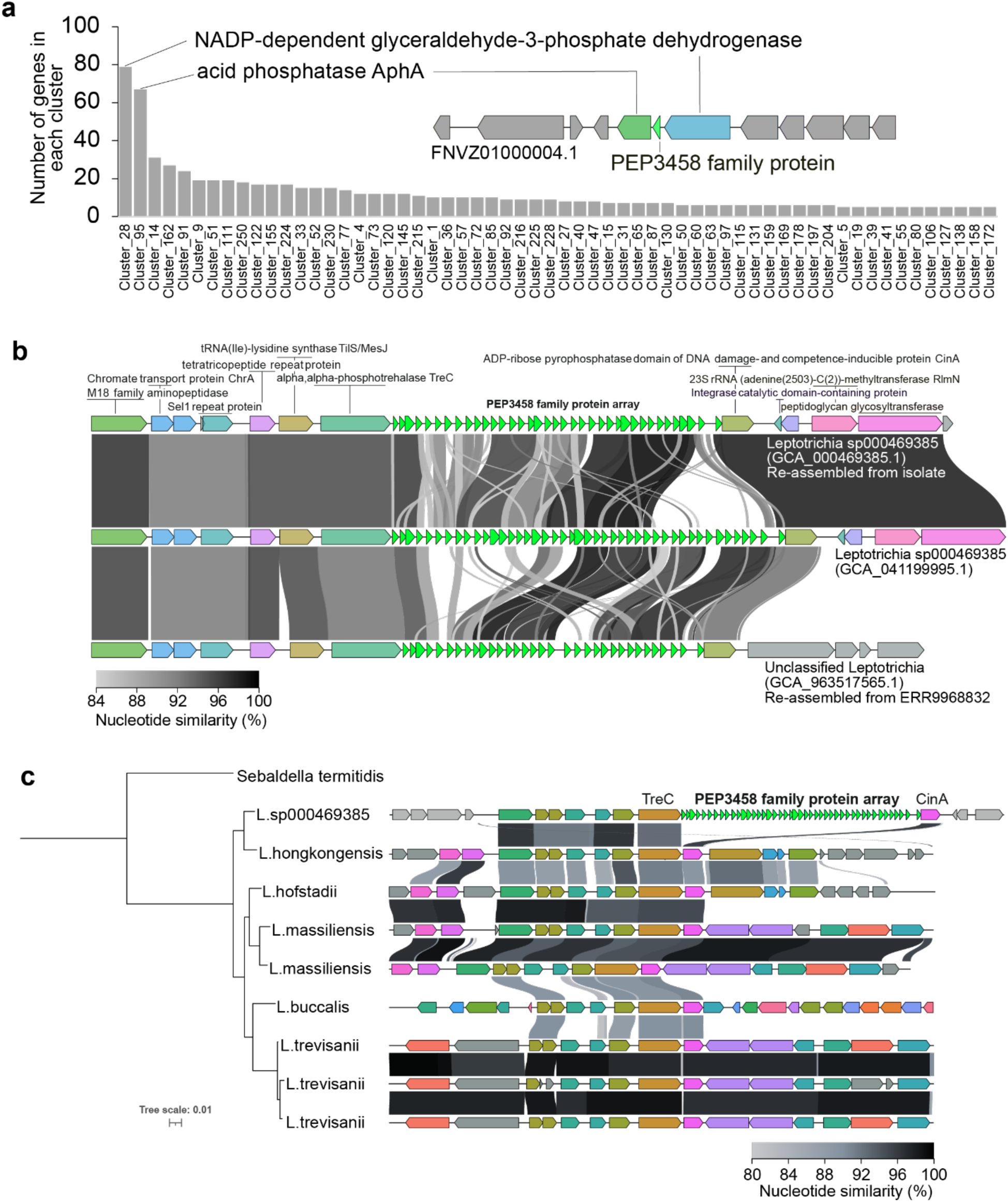
Genomic neighborhood and synteny of PEP3458 family protein arrays. a. Predicted functions of genes in the PEP3458 family gene neighborhood. Genes located one position upstream and downstream of each PEP3458 family ORF and protein array were extracted and clustered using CD-HIT with >90% identity. The x-axis represents the cluster ID; the y-axis shows the number of genes in each cluster. The predicted functions of the two largest clusters are annotated. The genetic map on the right shows a representative genomic locus (FNVZ01000004.1) encoding a Cluster_28 gene (NADP-dependent glyceraldehyde-3- phosphate dehydrogenase, green), a PEP3458 family ORF (green), and a Cluster_95 gene (acid phosphatase *aphA*, blue). b. Synteny of the PEP3458 array locus across three *Leptotrichia* genomes. Synteny comparison of the PEP3458 family protein array locus across three *Leptotrichia* genomes: *Leptotrichia* sp000469385 (GCA_000469385.1, re-assembled from an isolate), *Leptotrichia* sp000469385 (GCA_041199995.1), and unclassified *Leptotrichia* (GCA_963517565.1, re- assembled from metagenome ERR9968832). PEP3458 family ORFs are shown as green arrows; selected flanking genes are colored and labeled by predicted function. The genes satisfying >40% identity are shown as the same color. Gray ribbons connect homologous regions, shaded by nucleotide similarity (%). c. Interspecies synteny and phylogeny of the PEP3458 array locus across *Leptotrichia* species. The phylogenetic tree was reconstructed with *Sebaldella termitidis* as the outgroup. The synteny plot shows genomic loci surrounding the PEP3458 family protein array; PEP3458 family ORFs are shown as green arrows, *treC* and *cinA* are labeled, and flanking genes are colored by orthologous groups. Gray ribbons connect homologous regions, shaded by nucleotide similarity.

**Extended Data 10.**
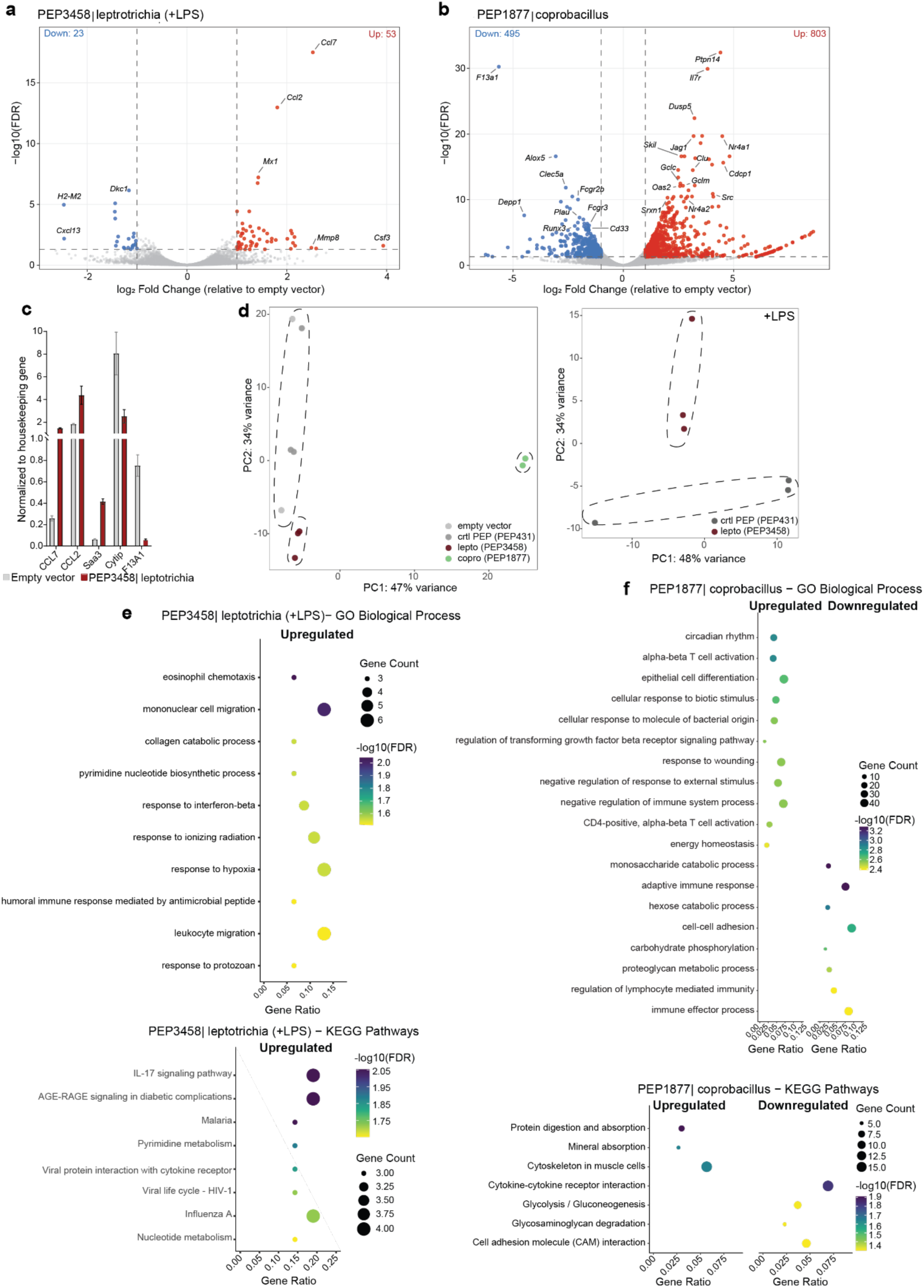
Bacterial microproteins induce distinct immune signatures in J774 macrophages. a. Volcano plot of *Leptotrichia* PEP3458 + LPS, showing DEGs, relative to control PEP431, with top DEGs are labeled (n=3 distinct biological replicates per condition). Differentially expressed genes were identified using DESeq2 with thresholds of |log2 fold change| > 1 and FDR < 0.05. Pathways sorted from top to bottom from highest to lower -log10(FDR) score b. Volcano plot of *Coprobacillus* PEP1877 vs Empty Vector showing DEGs, relative to control PEP431, with top DEGs labeled (n=2 distinct biological replicates of Coprobacillus 1877, n=3 distinct replicates of empty vector control), with the same thresholds as previously described. c. qRT-PCR of candidate genes identified in bulk RNAseq assay from J774 macrophages expressing either empty vector or *Leptotrichia* PEP3458. d. Over Representation Analysis (ORA) of Leptotrichia PEP3458 + LPS DEGs. Left: GO Biological Process enrichment showing upregulation of immune response pathways and metabolic processes. Right: KEGG pathway enrichment highlighting IL-17 signaling, viral infection response pathways, and nucleotide metabolism. Dot size represents gene count. Color intensity indicates statistical significance (-log10 FDR); upregulated pathways are shown in top panels/right side, downregulated in bottom panels/left side. Redundant GO terms were filtered using hierarchy-based semantic similarity (see Methods). FDR, false discovery rate (Benjamini-Hochberg correction). e. ORA of *Coprobacillus* PEP1877 DEGs. Top: GO Biological Process showing upregulation of T cell differentiation and circadian rhythm pathways. Bottom: KEGG pathways showing upregulation of TNF signaling and viral response pathways, with downregulation of metabolic pathways.

**Extended Data 11.**
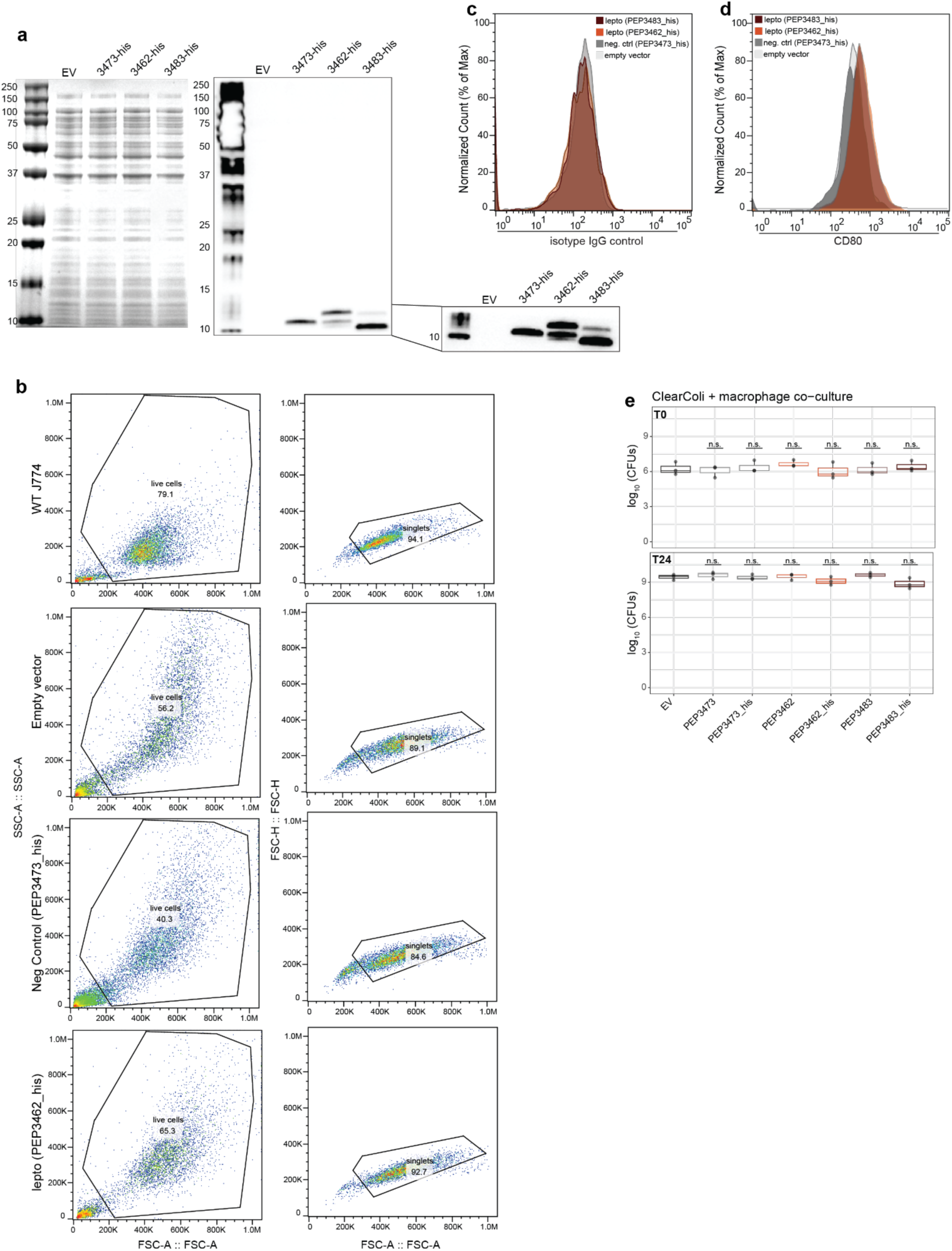
Infection of J774 macrophages with ClearColi expressing *Leptotrichia* microproteins induced an enhanced inflammatory response. a. Coomassie stain (left) and western blot (right) confirming expression of *Leptotrichia* microproteins (PEP3462 and PEP3483) and control microprotein, PEP3473. The zoomed image shows the blot after exposure without the signal saturation of the upper ladder bands. b. Flow cytometry plots demonstrating similar cell characteristics between J774 macrophages exposed to ClearColi strains expressing different microprotein (PEP3473, PEP3462, and PEP3483). c. Flow cytometry histogram plots demonstrating increased CD80 expression of J774 macrophages co-cultured with ClearColi expressing non-his tagged *Leptotrichia* microproteins (PEP3462, orange and PEP3483, red), relative to negative controls (PEP3473, dark grey and empty vector, light grey) d. Flow cytometry histogram plots demonstrating equivalent signal from isotype IgG control staining of J774 macrophages co-cultured with ClearColi expressing *Leptotrichia* microproteins (PEP3462, orange and PEP3483, red) and negative controls (PEP3473, dark grey and empty vector, light grey) e. Colony forming units of indicated ClearColi strains (empty vector, PEP3473, PEP3462, and PEP3483, with and without his tags) demonstrating equivalent bacterial counts at both the time of incubating with macrophages as well as upon harvesting for flow cytometry and RNAseq. n= 3 distinct biological replicates for each condition. Statistical testing performed with Wilcoxon signed-rank test.

**Extended Data 12.**
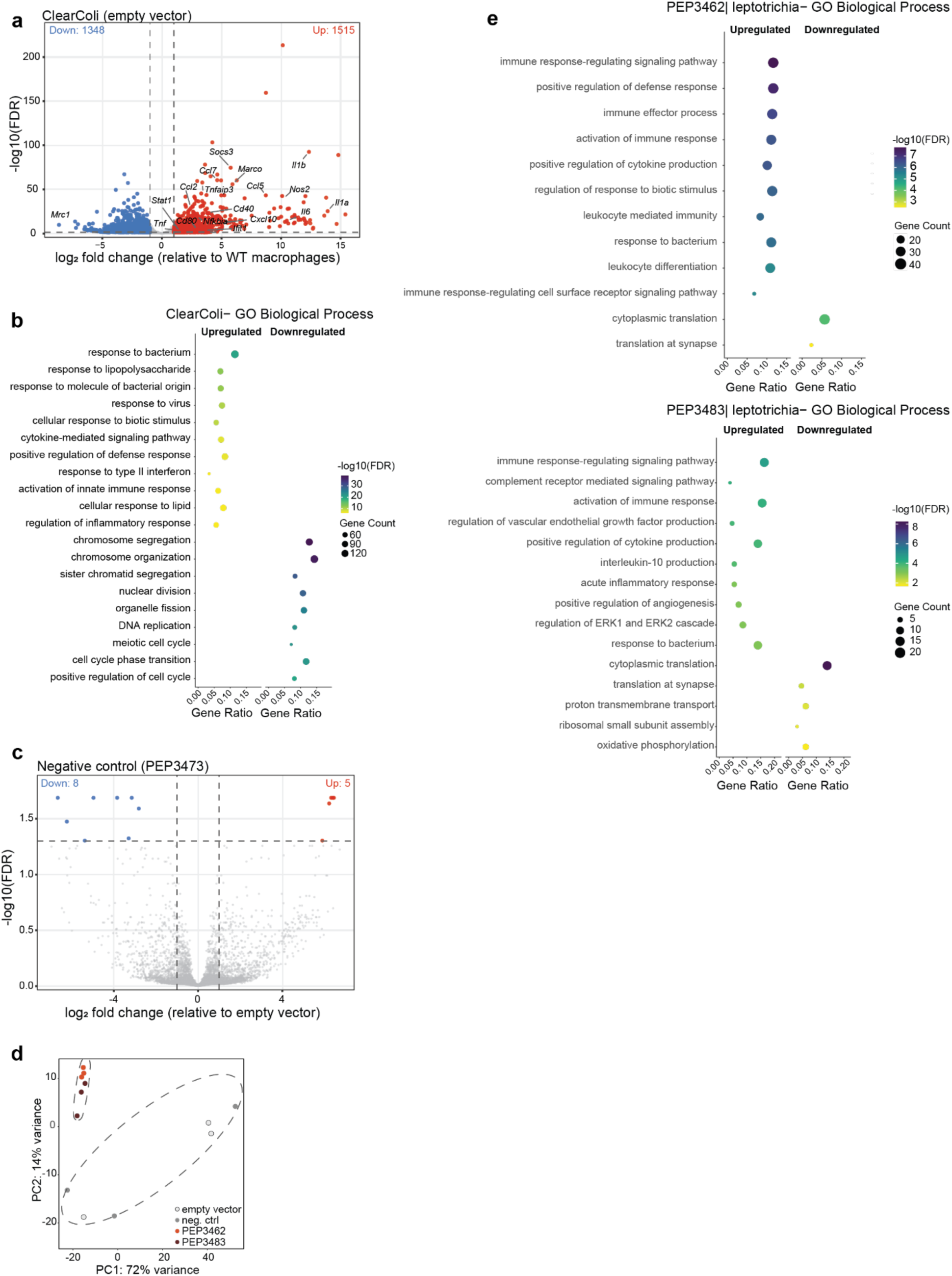
Increased inflammatory transcriptomic signatures in J774 macrophages co-cultured with ClearColi expressing *Leptotrichia* microproteins. a. RNA-seq demonstrating increased expression of J774 macrophages exposed to ClearColi (empty vector strain), relative to WT conditions. n=3 distinct biological replicates for all samples. Volcano plot shows DEGs, with red dots indicating upregulated genes and blue dots indicating downregulated genes. M1 associated, pro-inflammatory genes labeled. b. Over Representation Analysis (ORA) of macrophages exposed to ClearColi. GO Biological Process enrichment shows upregulation of immune response pathways and response to infection. Dot size represents gene count. Color intensity indicates statistical significance (- log10 FDR); upregulated pathways are shown in top panels/right side, downregulated in bottom panels/left side. Redundant GO terms were filtered using hierarchy-based semantic similarity, and manually pruned if further redundancy was present. c. RNA-seq demonstrating minimal differences between J774 macrophages exposed to ClearColi strains expressing PEP3473, relative to the empty vector strains. Volcano plot shows DEGs, with red dots indicating upregulated genes and blue dots indicating downregulated genes. d. Principal component analysis plots of bulk RNA-seq demonstrating clustering of macrophages expressing *Leptotrichia* microproteins, PEP3462 (orange) and PEP3483 (red), relative to relative to negative controls (negative control PEP3473 and empty vector, in dark and light grey). e. Over Representation Analysis (ORA) demonstrating enrichment of immune responses for macrophages co-culture with ClearColi expressing *Leptotrichia* microproteins PEP3462 (top) and PEP3483 (bottom). GO Biological Process enrichment shows upregulation of immune response pathways and response to infection. Dot size represents gene count. Color intensity indicates statistical significance (-log10 FDR); upregulated pathways are shown in top panels/right side, downregulated in bottom panels/left side. Redundant GO terms were filtered using hierarchy-based semantic similarity, and manually pruned if further redundancy was present.

## Supplementary Tables

Table 1. A description of the microproteins synthesized to test for immunomodulatory activity.

Table 2. DEseq analysis from cytokine display system

Table 3. GSEA from cytokine display experiments

Table 4. Processed counts and log fold enrichment scores from microprotein screen

Table 5. *Leptotrichia* microprotein clustering analysis

Table 6. DEseq analysis from microproteins hits expressed in peptide display system

Table 7. ORA of microproteins hits expressed in peptide display system

Table 8. DEseq analysis from ClearColi and macrophage co-culture experiments

Table 9. ORA from ClearColi and macrophage co-culture experiments

Table 10. Bacterial plasmids and strains

## Acknowledgements

We would like to thank members of the Bassik lab for their assistance with experimental design and computational analysis, including Peter Du and Nora Enright. We would also like to thank Jennifer Wargo and Nadim Ajami for helpful discussions, and Jessica Mark Welch (Forsyth Institute) for assistance in acquiring *Leptotrichia* isolates. This work used supercomputing resources provided by the Stanford Genetics Bioinformatics Service Center, supported by The project described was supported in part by ARRA (award no. 1S10RR026780-01) from the National Center for Research Resources. This work was supported by NIH R01CA301727 and a Paul Allen Investigator Award to Bhatt and Bassik labs. M.N.R is supported by Award Number K12-HD000850 from the Eunice Kennedy Shriver National Institute of Child Health and Human Development. Y.K. is supported by Stanford Medicine Children’s Health Center for the IBD and Celiac Disease postdoctoral fellowship. The Bhatt lab is supported by a philanthropic gift from Yosemite, a Stand up 2 cancer convergence grant, NIH U54AG089334, NIH U01DE035635 and NHLBI R01HL181969. The Bassik lab is supported by NHGRI R01HG011866 and NCI U54CA261719.

## Author Contributions

A.S.B and M.C.B conceived of and designed the study. M.N.R, J.L., Y.K., F.T.H., L.D. performed experiments, and M.N.R. performed analysis of experimental data sets. M.C, M.P.G, D.M., B.D., A.L conceptualized and designed the microprotein library. K.S performed library cloning. Y. K. performed computational analysis related to microprotein arrays. The manuscript was drafted and all figures prepared by M.N.R. and Y.K. The manuscript was finalized by M.N.R., Y.K., L.B., A.S.B and M.C.B.

